# Overlaid positive and negative feedback loops shape dynamical properties of PhoPQ two-component system

**DOI:** 10.1101/2020.07.09.194944

**Authors:** Satyajit D Rao, Oleg A Igoshin

**Affiliations:** Department of Bioengineering, Rice University, Houston, TX, USA

## Abstract

Bacteria use two-component systems (TCSs) to sense environmental conditions and change gene expression to adapt to those conditions. To amplify cellular responses, many bacterial TCSs are under positive feedback control, i.e. increase their own expression when activated. In *E. coli*, Mg^2+^-sensing TCS, PhoPQ, in addition to the positive feedback includes a negative feedback via upregulation of MgrB protein that inhibits PhoQ. How interplay of these feedback loops shapes steady state and dynamical responses of PhoPQ TCS to change in Mg^2+^remains poorly understood. In particular, how the presence of MgrB feedback affects the robustness of PhoPQ response to overexpression of TCS is unclear. It is also unclear why the steady state response to decreasing Mg^2+^is biphasic, i.e. plateaus over a range of Mg^2+^concentrations and then increases again at growth-limiting Mg^2+^. In this study, we use mathematical modeling to identify potential mechanisms behind these experimentally observed dynamical properties. The results make experimentally testable predictions for the regime with response robustness and propose novel explanation of biphasic response constraining the mechanisms for modulation of PhoQ activity by Mg^2+^and MgrB. Finally, we show how interplay of positive and negative feedback loops affect networks steady-state sensitivity and response dynamics. In the absence of MgrB feedback, the model predicts oscillations thereby suggesting a general mechanism of oscillatory or pulsatile dynamics in autoregulated TCSs. These results help better understanding of TCS signaling and other networks with overlaid positive and negative feedback.

**Author summary:** Feedback loops are commonly observed in bacterial gene-regulatory networks to enable proper dynamical responses to stimuli. Positive feedback loops often amplify the response to stimulus, whereas negative feedback loops are known to speed-up the response and increase robustness. Here we demonstrate how combination of positive and negative feedback in network sensing extracellular ion concentrations affects its steady state and dynamic responses. We utilize published experimental data to calibrate mathematical models of the gene regulatory network. The resulting model quantitatively matches experimentally observed behavior and can make predictions on the mechanism of negative feedback control. Our results show the advantages of such a combination feedback loops and predict the effect of their perturbation on the steady state and dynamic responses. This study improves our understanding of how feedback loops shape dynamical properties of signaling networks.

## Introduction

Bacteria use two component systems (TCSs) to sense and respond to environmental stimuli [1,2]. TCSs are also widely used in synthetic biology applications to sense specific stimuli and control gene expression [3–5]. A TCS consists of a sensor kinase often located on the inner membrane and a cognate response regulator protein located in the cytoplasm. The sensor kinase senses environmental stimulus and responds by autophosphorylating at the histidine residue [6]. Phosphorylated kinase catalyzes a transfer of phosphate to the response regulator. In absence of environmental stimuli, sensor kinases sometimes have phosphatase activity, i.e. they can dephosphorylate the response regulator. The phosphorylated response regulator is transcriptionally active, initiating cellular response. As part of cellular response, the response regulator often activates transcription of genes encoding the two components themselves [7], creating a positive feedback loop.

The Mg^2+^-sensing PhoPQ TCS is found in many bacterial species, such as *Salmonella, Yersinia pestis* and *E. coli* [8–14]. The sensor kinase PhoQ responds to low extracytoplasmic Mg^2+^levels, acidic pH and antimicrobial peptides. In high Mg^2+^, the periplasmic sensing domain of PhoQ is bound to Mg^2+^resulting in a conformation of PhoQ that has low autokinase activity but high phosphatase activity towards phosphorylated PhoP (PhoP-P) [8]. That keeps the expression of PhoP-P-dependent genes low. In response to Mg^2+^limitation, dissociation of Mg^2+^from PhoQ promotes a conformational change that increases the autokinase activity and suppresses the phosphatase activity [15]. That leads to accumulation of PhoP-P and increase in the expression of its regulon. PhoP-P regulons vary significantly between different bacterial species but retain a few common features. First, the PhoPQ TCS upregulates transcription of its own operon *phoPQ*. This upregulation leads to a positive feedback in the system. Second, PhoP activates transcription of a small lipoprotein (SlyB in *Salmonella* and *Yersinia pestis*, MgrB in *E. coli*) that limits kinase activity (Fig 1) [16–18]. These interactions form a negative feedback loop.

**Fig 1.**
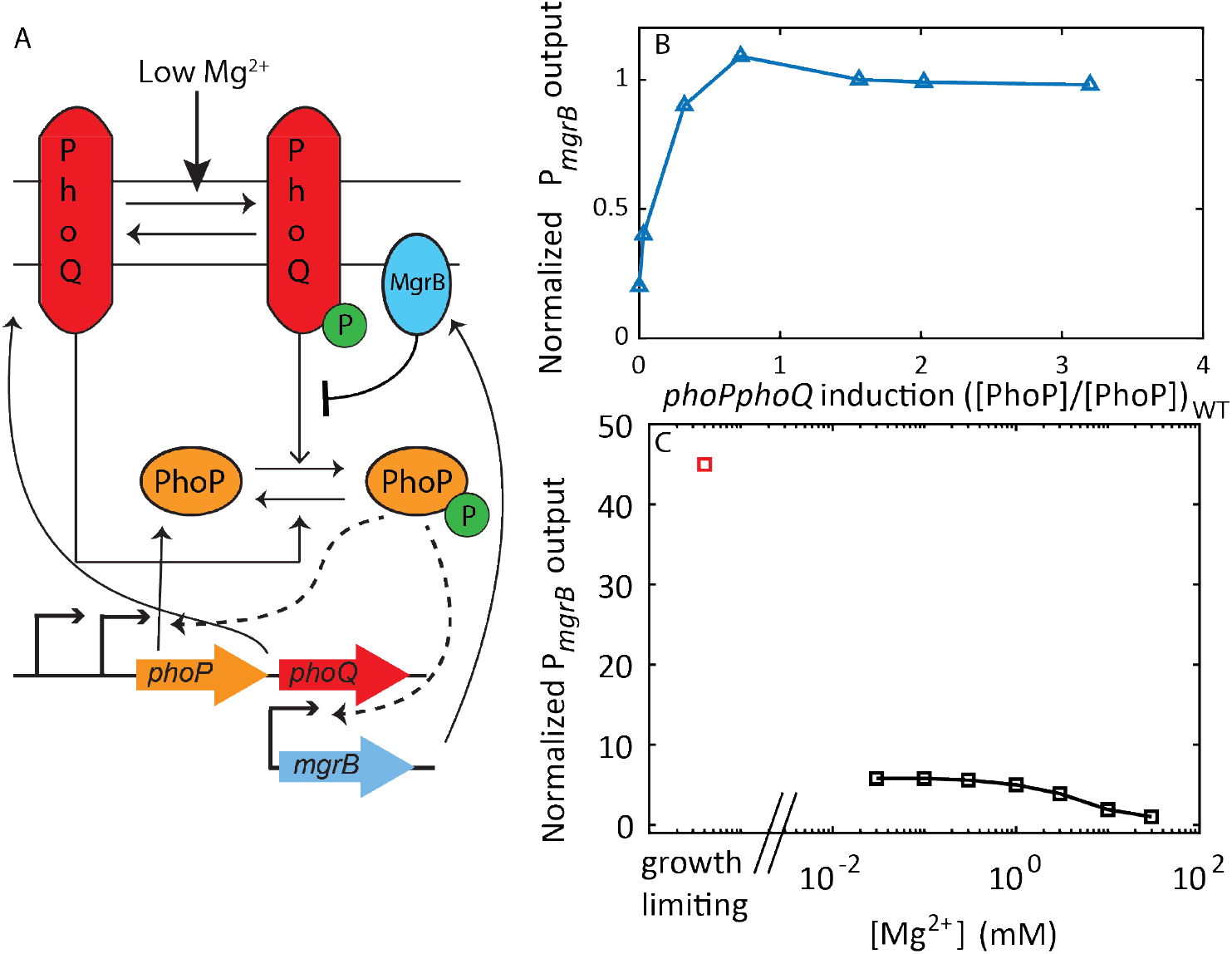
PhoPQ two component system senses low Mg^2+^ concentrations through direct interactions with PhoQ. (A) When activated by low external Mg^2+^PhoQ undergoes autophosphorylation, transfers phosphoryl group to PhoP which activates transcription of downstream genes. PhoP positively regulates transcription of *phoPQ* operon, as well as mgrB. MgrB binds PhoQ and suppresses kinase activity. (B) Normalized reporter output from P_*mgrB*_ saturates as *phoPQ* operon transcription is increased. (C) Steady state normalized reporter output (P_*mgrB*_) plateaus as Mg^2+^ decreases, but increases further at growth limiting conditions (hypothetical normalized reporter output at growth limiting Mg^2+^, red square). Plot recreated from [19]

How does positive autoregulation of PhoP/PhoQ affect phosphorylation level of PhoP? Notably, over a range of low Mg^2+^concentrations, elimination of autoregulation of PhoPQ results in no significant difference in the PhoP-P activity measured via transcriptional reporter of the *mgrB* promoter [19]. This observation suggests that PhoP-P level is insensitive (robust) to increase of *phoPQ* operon production. This robustness was confirmed by monitoring PhoP activity reporter in a strain with a chemically inducible *phoPQ* promoter (Fig 1); increasing PhoPQ expression post wild-type level does not change the reporter level [19]. Previously published mathematical models show that this robustness arises due to bifunctionality of the kinase [19–21]. However, these models may not be directly applicable for *E. coli* PhoPQ TCS as they do not account for PhoPQ interactions with MgrB protein. While the exact mechanism by which MgrB modulates PhoQ activity is unknown, MgrB specifically inhibits kinase activity through direct interaction [17]. A strain lacking both the *mgrB* gene and PhoQ-phosphatase activity shows higher promoter activity compared to a strain merely lacking the PhoQ-phosphatase activity [16]. Since overexpressing PhoQ could in-principle outcompete the inhibitory effect of MgrB, understanding robustness of PhoP-P to PhoPQ overexpression requires models that explicitly include negative feedback regulation.

Notably, robustness of PhoP activity to elimination of PhoPQ autoregualtion is no longer observed in growth-limiting Mg^2+^levels (< 10^−3^ mM) [19]. Furthermore, in these conditions, the PhoP activity greatly exceeds the activity observed over a range of low but not growth-limiting (between 1 and 0.01 mM) Mg^2+^ concentrations. Notably: PhoP activity is nearly the same over that range of Mg^2+^levels forming a plateau between 1 and 0.01 mM Mg^2+^following a gradual increase from 100 to 1 mM Mg^2+^. Such a plateau has been observed for multiple promoters with varying affinities to PhoP-P, suggesting this pattern is not a property of one particular promoter [9]. Interestingly, at 0.01 mM Mg^2+^the levels of PhoP-P are such that the promoters remain far from saturation [9].

A combination of plateau in promoter activity over a range of low Mg^2+^levels with further increased activity in growth-limiting Mg^2+^can be referred to as biphasic dose-response (Fig 1). Miyashiro and Goulian hypothesize that this biphasic dose-response is indicative of Mg^2+^binding to PhoQ at multiple sites with different affinities [9]. However, this hypothesis has not been tested experimentally or theoretically. Alternatively, feedback architecture might shape a biphasic dose-response. If negative feedback dominates over a range of low Mg^2+^concentrations, while positive autoregulation is strongly activated only in growth-limiting Mg^2+^, we could perhaps expect steady state PhoP-P to display two phases of activation. Since it is unclear how overlaid positive and negative feedback loops shape observed dose-response, detailed mathematical models of PhoPQ TCS can be used to understand steady state PhoP-P as a function of Mg^2+^.

In this study, we use mathematical modeling to understand how positive and negative feedback loops interact to shape dynamical properties of the PhoPQ TCS in *E. coli*. First, we identify conditions under which PhoP-P remains robust to *phoPQ* overexpression even in presence of MgrB-mediated negative feedback. Next, we search for mechanisms underlying the biphasic dose-response. We use published temporal and steady state data for wild type and mutant *E. coli* strains to calibrate our models. Finally, we use these calibrated models to understand advantages of the overlaid positive and negative feedback design of the PhoPQ system. Taken together, this study shows how mathematical modeling and experimental data can be used together to understand the relationship between network structure and cellular function in bacteria.

## Results

### PhoPQ TCS can show robustness if MgrB is in excess of PhoQ

Structural sources of robustness to variation in species concentrations have been identified previously for mass-action reaction networks [21, 22]. Two component systems (TCSs) with bifunctional kinases are known examples of such biochemical networks. The concentration of phosphorylated response regulator can be robust to changes in total concentrations of sensory kinase and response regulator proteins [20–22]. To ascertain if the biochemical reaction network of PhoPQ TCS meets the criteria for absolute concentration robustness (ACR) put forth by Shinar and Feinberg [22], we analyze the reaction network with or without MgrB (Figure 1 in S1 Text). While a reaction network without MgrB did in fact meet the criteria to obtain ACR, the reaction network of PhoPQ with MgrB network did not (Table 1 in S1 Text). This analysis suggests that in contrast to a typical TCS featuring a bifunctional kinase, robustness to total protein concentrations is not theoretically predicted by the structure of the reaction network. To understand that result we note that ACR occurs due to ability of PhoQ to control both phosphorylation and dephosphorylation flux to PhoP. Increase in PhoQ level will increase both fluxes proportionally without affecting PhoP-P. On the other hand, when MgrB-mediated inhibition of kinase activity is present, increase in PhoQ concentration disproportionally increases dephosphorylation flux.

To find conditions under which PhoP-P could be robust to total-PhoPQ expression, we modify the PhoPQ model in ref [19] to explicitly include negative feedback regulation. We consider a system without positive feedback, i.e. with total-PhoPQ expression controlled independent of PhoP-P. To simplify steady state analysis of PhoP-P, we follow the approach used in ref [19] and break the model into two modules (Fig 2A). For the transcription module the input is PhoP-P and the output is total-MgrB. We use a standard Hill-function to describe how transcription rate of *mgrB* and correspondingly total MgrB concentration depends [PhoP-P](Eq. 1).

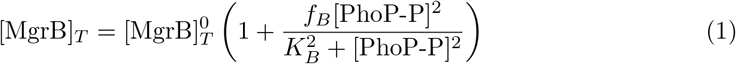

**Fig 2.**
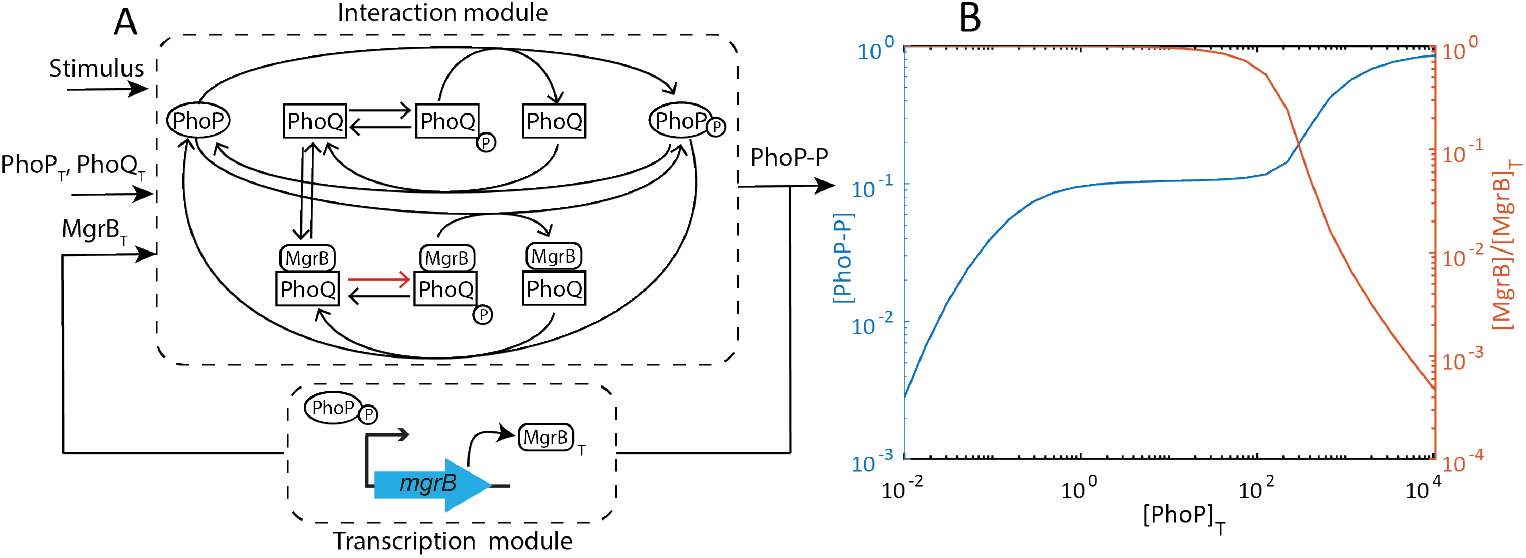
PhoP-P is robust to overexpression of PhoP, PhoQ. A – Modified PhoPQ TCS interaction network shows PhoQ binding MgrB, repressing PhoQ autophosphorylation. This forms the interaction module. The interaction module takes 3 inputs (i) Stimulus (autophosphorylation rate), (ii) Total PhoP (PhoQ is assumed proportional, and 1/40 times PhoP based on actual measaurements) and (iii) Total MgrB. The system is numerically solved for a steady state concentration of PhoP-P as a function of varying PhoP, PhoQ total. The interaction module is coupled with a transcription module representing negative feedback with PhoP-P as input and total MgrB concentration as output. B – The system is solved numerically for steady state concentration of PhoP-P as a function of varying PhoP (and PhoQ) total (blue). PhoP-P is robust to PhoP/PhoQ concentrations, increasing further when PhoQ concentration is large enough to overcome MgrB negative feedback. Over most of the range of PhoQ concentrations, MgrB ≈ MgrB-total indicating large stoichiometric excess MgrB (orange line). Robustness breaks when MgrB is no longer in large excess of PhoQ

Here 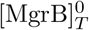 denotes minimum MgrB concentration at basal expression of *mgrB, f_B_* represents maximal fold upregulation of MgrB while *K_B_* denotes half activation concentration of PhoP-P. The interaction module consists of the cycle of phosphorylation-dephosphorylation catalyzed by PhoQ and PhoQ-MgrB. For the interaction module the inputs are total-PhoP, PhoQ and MgrB as well as the stimulus level (i.e. autophosphorylation rate of PhoQ). Steady state is at the intersection of the two modules (S1 Fig). As in the model in ref [19], we assume total-PhoP/PhoQ ratio is constant, and this ratio is greater than 1 [23]. To model MgrB inibition, we assume that when MgrB binds to PhoQ, its autophosphorylation rate (Fig 2A) decreases by a factor λ < 1. The remaining rate constant parameters are assumed same for PhoQ and PhoQ-MgrB states.

With this model we investigate how steady state [PhoP-P] depends on total PhoP, PhoQ (see S2 Text for full analysis). Basal expression of MgrB (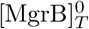 in Eq. 1) is a free parameter. For each value of 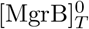 we can solve the two modules for a range of total-PhoP, PhoQ values. We find that steady state [PhoP-P] is not in general robust (S1 Fig). However, at high 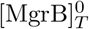 values, PhoP-P can be robust over a limited range of total-PhoP (Fig 2B).

To understand how steady state PhoP-P can show robustness to only a limited range of total-PhoP, and only at high 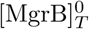 values, we look at phosphorylation and dephosphorylation fluxes of PhoP (S2 Text). Equating phosphorylation and dephosphorylation fluxes, we can obtain an expression for [PhoP-P] shown in Eq. 2

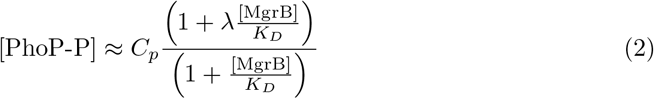

In this equation, [PhoP-P] depends on concentration of free MgrB. Since concentration of free MgrB depends on concentration of PhoQ in general, there is no robustness. However, when MgrB is in large excess of PhoQ, the function simplifies to Eq. 3.

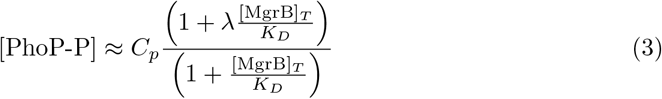

Here *C_p_* is a combination of parameters as noted previously in refs [19,20], and *K_D_* is the dissociation constant for [PhoQ-MgrB]. In this expression, [PhoP-P] then depends only on MgrB-total, which in turn depends on [PhoP-P] (Eq. 1). Thus, steady state [PhoP-P] remains independent of total PhoQ, PhoP. Indeed, Output ceases to be robust once [PhoQ]_*T*_ ~ [MgrB]_*T*_. This is illustrated by a plot of [MgrB]/[MgrB]_*T*_ (Fig 2B). Thus our model of PhoPQ-MgrB with negative feedback shows that robustness to total-PhoP/PhoQ is not due to the cycle of phosphorylation alone, but can be obtained if MgrB is much more abundant than the kinase PhoQ.

### Models with autophosphorylation suppression by MgrB alone cannot explain biphasic dose-response

Given that the previous model of autophosphorylation suppression by MgrB can explain robustness of PhoP-P to total-PhoPQ levels, we explore steady state dose-response behavior of the model. Specifically, we investigate whether the model can recreate a biphasic dose-response, i.e. show an intermediate plateau (Fig 1C). To compare with experimental data of ref [19], we construct a detailed dynamic model with two reporter proteins YFP and CFP [9] (S1 Appendix). To calibrate the model, we fit simulated values of YFP:CFP to reported values from various experiments (Methods). Time-course measurements in wild-type cells switched from high to low Mg^2+^levels (published in [16]) were used to fine tune temporal parameters. The Mg^2+^step down experiment was also conducted with several mutant strains. Measurements from these mutant strains can serve as important biological constraints on the model. Thus, we simulate YFP:CFP values with in-silico mutants and fit to respective experimental values (Methods).

Steady state values of YFP:CFP over a range of Mg^2+^concentrations have also been measured for wild-type cells (published in [19]). We use these measurements to tune steady state parameters. In addition, we introduced a qualitative condition for greater reporter output at very high stimulus to recapitulate effects at growth limiting Mg^2+^ [19]. Normalized experimental data used, simulation protocols and parameter fitting procedure are described in Methods.

Over multiple parameter fitting attempts, the model could only show graded increase followed by a plateau in steady state promoter activity (S3 Fig). Thus, we hypothesize that models of PhoPQ-MgrB with more complex mechanisms are required to explain biphasic dose-response.

### Models representing PhoQ with separate kinase and phosphatase conformations can explain biphasic dose-response

To explain biphasic dose-response, we constructed a model of PhoQ with an explicit Mg^2+^sensing mechanism (S1 Appendix). While understanding of how Mg^2+^modulates PhoQ activity is still incomplete in *E. coli*, research in *Salmonella* has suggested that a conformation change resulting from Mg^2+^binding to PhoQ increases phosphatase activity [8]. Based on this finding we hypothesize two conformations of PhoQ: phosphatase (PhoQ) and kinase (PhoQ*) (Fig 3A). Extracellular Mg^2+^binds to PhoQ* and drives a transition to PhoQ thus shutting off PhoP-P activity. We assume that extracellular Mg^2+^concentration does not change over time and include it in the rate constant of switching from PhoQ* to PhoQ (Fig 3A) i.e. 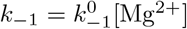. For simplicity we assume that only the kinase conformation undergoes autophosphorylation and phosphotransfer steps, while only phosphatase conformation dephosphorylates PhoP-P. MgrB can bind both conformations of PhoQ independently, subsequently modulating one or more rates.

**Fig 3.**
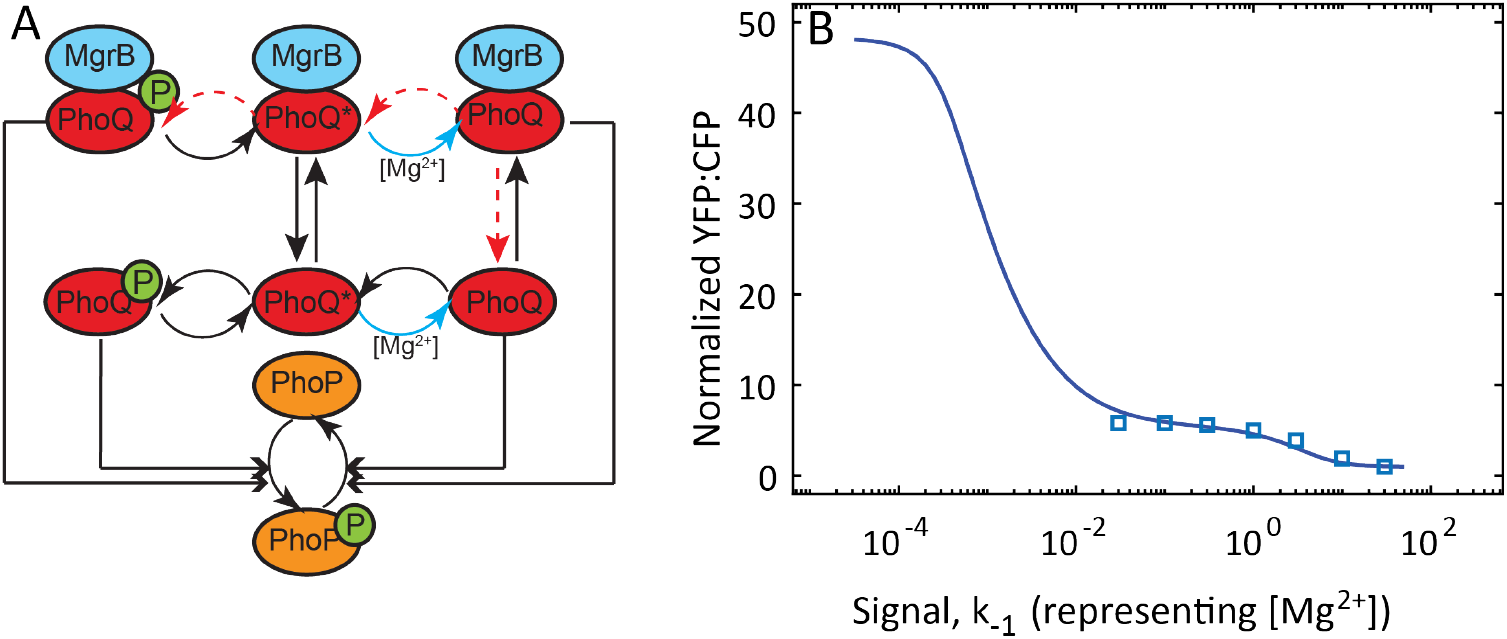
A two-state model of PhoPQ TCS can explain biphasic response. A-Schematic of the two-state model. PhoQ exists in phosphatase (PhoQ) or kinase (PhoQ*) form, PhoQ* assumed to bind Mg^2+^and switch to PhoQ. Concentration of Mg^2+^in medium assumed constant, and absorbed into a pseudo-first order kinetic rate, *k*_−1_ (blue arrow). MgrB reversibly binds PhoQ/PhoQ*. B – Simulated output (normalized YFP:CFP; Methods) from the ODE model representing schematic in A with two rate constants suppressed in MgrB bound PhoQ. The affected rates are denoted by red arrows: switching rate from phosphatase to kinase (i.e. *k*_1_, PhoQ-Mg^2+^dissociation), and autophosphorylation. Detailed balance condition is satisfied by assuming PhoQ-MgrB dissociation is suppressed by the same factor as *k*_1_. Simulated steady state output shows biphasic response to increasing signal.

The mechanism for MgrB mediated suppression of PhoQ remains unknown. Thus, we sought to understand which combination of rates of the phosphorylation cycle is likely to be modulated by MgrB. To this end, we implemented multiple models with MgrB affecting different rate constants in each. With each model we simulate a time course of YFP:CFP following downshifts in Mg^2+^for all the pairs of downshifts reported in Salazar et al [16] (S2 Fig). Time course simulations are performed for wild-type and in-silico mutants (S1 Appendix). In addition, we perform steady state dose-response simulations. We then obtain parameters that generate close fit with experimental data (Methods). We verify the accuracy of these parameters by performing Mg^2+^downshift simulations with an *in-silico* mutant expressing *mgrB* constitutively, as well as a PhoQ-phosphatase activity lacking mutant (S5 Fig). If simulations qualitatively match experimental data for the two mutant strains, we consider those parameters for further analysis. In dose-response simulations, we look for a second phase of strong promoter activation at very low Mg^2+^ (10^−4^ mM).

We find the biphasic dose-response pattern (Fig 3B) and closest matches with all experimental data (S4 Fig) in models where MgrB suppresses two rates - (i) autophosphorylation and (ii) activation (PhoQ → PhoQ* transition; red arrows, Fig 3A). Simulated steady state reporter output as a function of signal (*k*_−1_) shows two distinct ranges of signal where output increases, separated by a plateau (Fig 3B). Further, this model is able to fit time-course data for wild-type and mutant strains as well (S4 Fig). Interestingly, models with any other combinations of rate constants modulated by MgrB were unable to reproduce this biphasic response to signal. Thus, our analysis isolates a potential mechanism for MgrB suppressing PhoQ kinase activity. Taken together, models with an explicit Mg^2+^sensing mechanism and where MgrB modulates PhoQ phosphatase to kinase transition and autophosphorylation rates can explain the biphasic dose-response.

### Abundance of MgrB, strong suppression and slow transitions between PhoQ states together can create plateau in signal response

While this model can explain biphasic signal response, the mechanism behind a plateau at intermediate signal levels is not fully clear. To understand how steady state [PhoP-P] is insensitive to signal (*k*_-1_) in our model (Fig 4 D), we simplify the model so that analytical solutions will be possible in different ranges of signal. Matching phosphorylation and dephosphorylation fluxes can then provide expressions for steady state PhoP-P (see S4 Text for complete analysis). Analyzing how these fluxes change as a function of signal can clarify the mechanism behind PhoP-P plateau at *k*_−1_ values corresponding to 1-0.01 mM Mg^2+^range (Fig 4 B-D).

**Fig 4.**
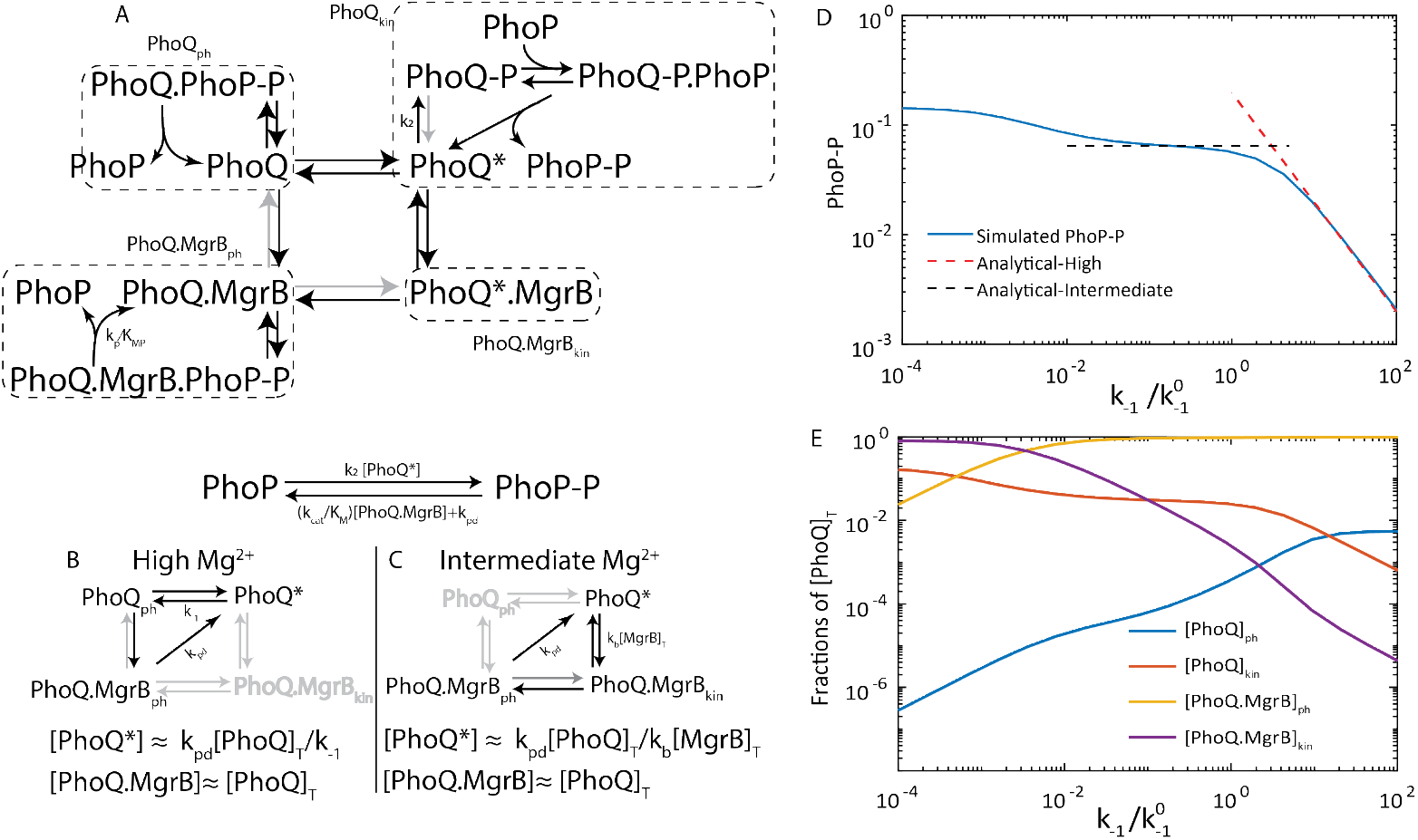
Understanding the biphasic dose-response possible in the two-state model of PhoPQ. A - Reactions in PhoPQ-MgrB network. Dotted squares enclose 4 sub forms of PhoQ (Q_ph_, Qkin, QB_ph_, QB_kin_). 4 reactions outside the dotted squares have rates comparable to dilution due to growth. Each sub form dilutes with a rate *k_pd_*, while synthesis is only in the Q* form. B - Most significant fluxes at high Mg^2+^. Steady state [PhoP-P] is approximated by matching phosphorylation and dephosphorylation fluxes. Phosphorylation flux is proportional to 1/*k*_−1_, while dephosphorylation flux is approximately constant. C-Most significant fluxes at intermediate Mg^2+^. Phosphorylation flux is proportional to [*B*]_*Total*_ and independent of *k*_−1_ while dephosphorylation flux is still approximately constant D-[PhoP-P] as a function of signal showing plateau at intermediate Mg^2+^. Blue line indicates simulated [PhoP-P] from the model in the previous section. Red dashed line shows approximate [PhoP-P] in the high Mg^2+^range, and black dashed line shows the approximate plateau value of [PhoP-P] in the intermediate Mg^2+^range. E-Fractions of total Q in the 4 catalytic forms.

Phosphorylation and dephosphorylation fluxes depend on the concentrations of the two catalytic states of PhoQ – kinase and phosphatase – and their MgrB-bound counterparts (Fig 4A). In our model we find that over high to intermediate Mg^2+^, nearly all PhoQ molecules exist in MgrB-bound phosphatase state (PhoQ.MgrB_ph_, Fig 4 E). We analyze steady state concentrations of PhoQ forms at the limit of complete suppression (S4 Text). We consider the limit in which MgrB binds PhoQ and completely suppresses autophosphorylation (Fig 4A) and activation rate constant (gray arrows Fig 4A). PhoQ.MgrB_ph_ concentration only reduces by dilution due to growth. In our model we find that over high to intermediate Mg^2+^, nearly all PhoQ molecules exist in MgrB-bound phosphatase state (PhoQ.MgrB_ph_, Fig 4E). Therefore, we neglect contribution of PhoQph to dephosphorylation of PhoP-P. Further, we assume that all the phosphate that enters the systems through autophosphorylation of PhoQ* transfers to PhoP (i.e. autodephosphorylation of PhoQ-P is negligible). The net phosphorylation flux of PhoP, then, is equal to autophosphorylation flux (proportional to PhoQ*). Dephosphorylation flux has two portions – phosphatase activity (proportional to [PhoQ.MgrB]), and dilution due to growth. The steady state [PhoP-P] is found by equating phosphorylation and dephosphorylation fluxes (S4 Text).

Thus, PhoP-P concentration depends on how [PhoQ*] and [PhoQ.MgrB] change as a function of *k*_−1_. This in turn depends on which fluxes dominate in signal ranges corresponding to high and intermediate Mg^2+^. We find that at high Mg^2+^, flux of [PhoQ*] deactivation (rate k_−1_) dominates over binding MgrB (rate *k_b_*[*MgrB*]; Fig 4B). In this range [PhoP-P] depends linearly on signal. An approximate analytical expression over a range of *k*_−1_ values corresponding to high Mg^2+^ matches simulated [PhoP-P] using parameters from the previous section (Fig 4D). Thus from high to intermediate Mg^2+^, the promoter output increases a few fold (Fig 3B). At intermediate Mg^2+^, however, flux of PhoQ* binding MgrB is much greater than deactivation (Fig 4C). In this range, [PhoP-P] depends on [*MgrB*]_*T*_ (which in turn depends on [PhoP-P]) but not signal, Fig 4 C). An analytical solution for a single steady state [PhoP-P]_*int*_ independent of signal can be found (Fig 4D). Taken together, strong suppression of kinetic rates by excess MgrB and growth dilution shape biphasic dose response of PhoPQ.

### Combination of positive and negative feedback increases range of sensitivity to signal

What advantages does this unusual combination of positive and negative feedback provide? We know from experimental observations that negative feedback creates partial adaptation and faster kinetics, while positive feedback amplifies output and helps cells survive in growth limiting magnesium. To find out how steady state behavior is shaped by overlapping feedback loops, we simulated steady state dose-response with only one feedback present at a time.

We find a narrow range of signal sensitivity with negative feedback absent, while a much wider range without positive feedback (Fig 5A and B). Thus, in addition to kinetic advantages, negative feedback keeps the system sensitive to changes in magnesium over a much wider range of concentrations. However, without positive feedback the maximum output is much lower than with both feedback loops present, validating experimental observations of strong stimuli activating positive feedback. Taken together we find that negative feedback allows the TCS to tune the competing activities of PhoQ with time (to create overshoot dynamics) as well as stimulus (biphasic dose-response).

**Fig 5.**
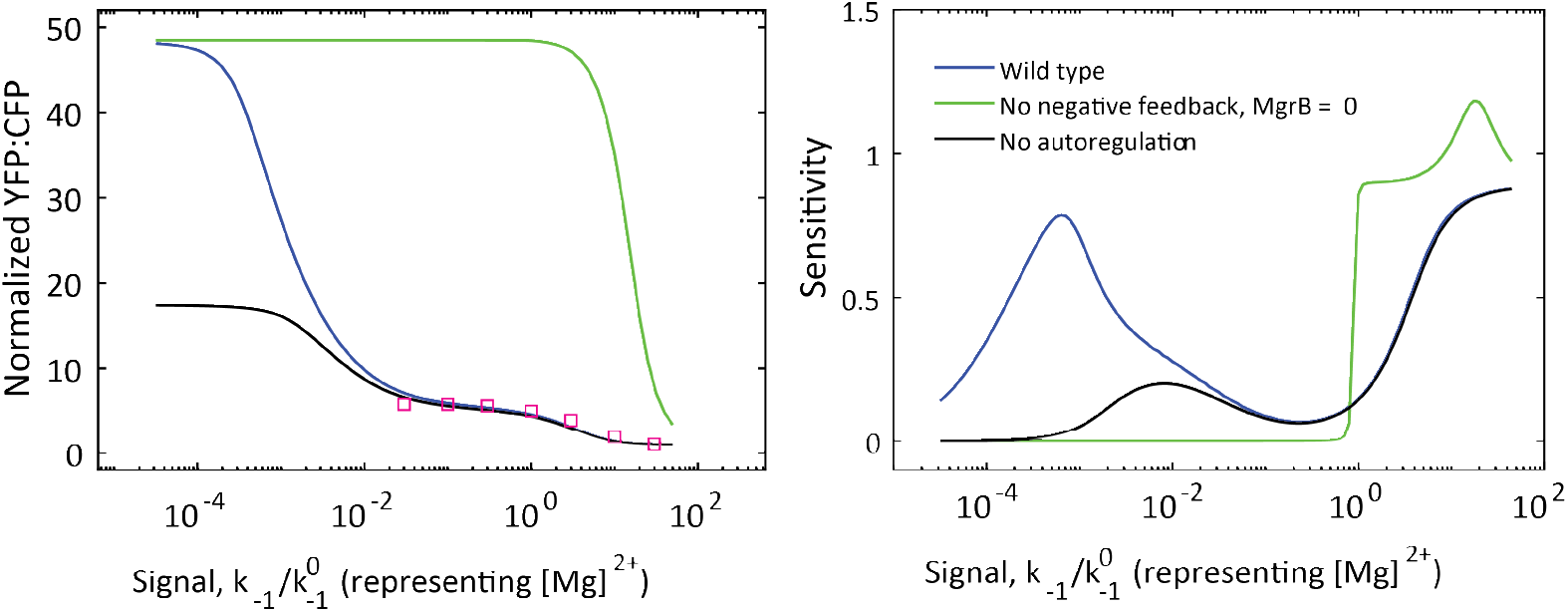
Combination of positive and negative feedback increases range of sensitivity to signal. A - steady state response of simulated promoter output (YFP, normalized to high Mg^2+^YFP) for models with no positive feedback (black), no negative feedback (green) and both feedback loops (blue). B-Absolute sensitivity (log derivative of PhoP-P with respect to *k*_−1_, 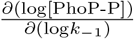) for model with no negative feedback (green) has the shortest range of signal sensitivity, while the wild-type model displays two phases of high sensitivity to signal.

### Negative feedback can suppress oscillations

In addition to increasing range of sensitivity to signal, we unexpectedly find that negative feedback through upregulation of *mgrB* may also prevent oscillations in the network. When investigating the dynamics of the responses for *in silico* mutants lacking negative feedback, i.e. with constitutive *mgrB* expression(Fig 6A), we discovered limit cycle oscillations are observed at intermediate signal levels. These oscillations are only seen for low *mgrB* expression rates, i.e. with MgrB level comparable to that in unstressed wild-type cells (Fig 6B). Notably, these oscillations are observed for all the parameter sets that fit experimental data and show a biphasic dose response for wild-type cells (Figure 1 in S3 Text). However, the oscillations were absent for the wild-type dynamics, i.e. when negative feedback is present(Fig 6B, blue line). The result is unexpected since oscillations require negative feedback and in our case elimination of negative feedback leads to oscillations.

**Fig 6.**
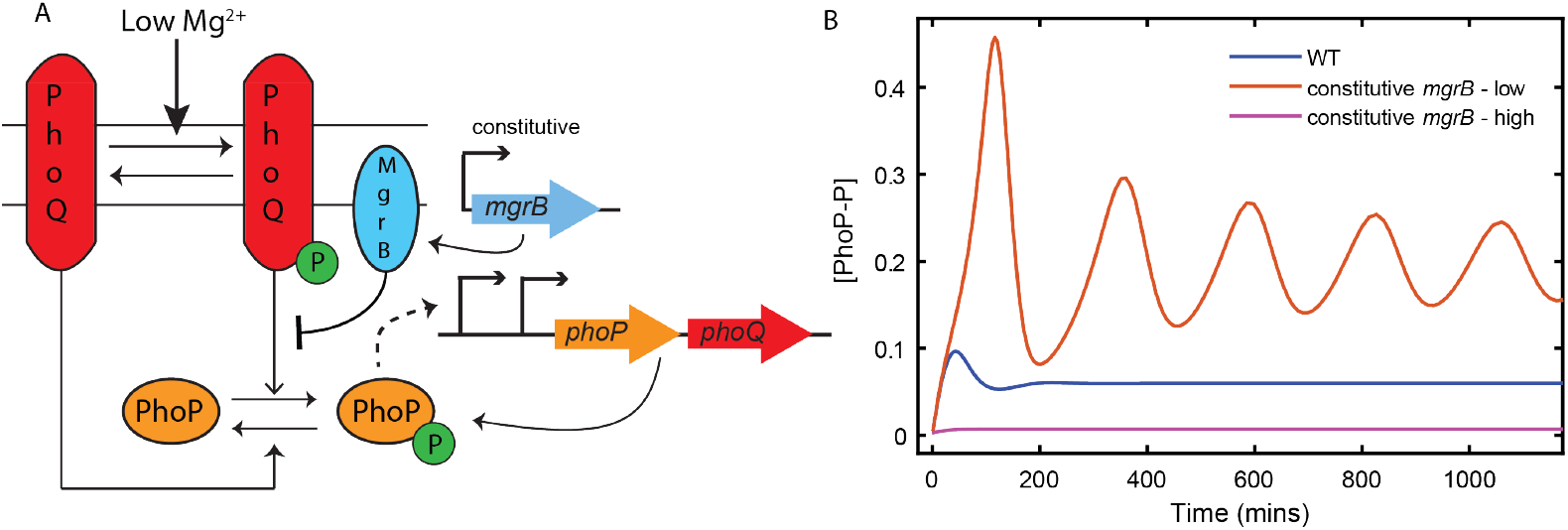
Removing upregulation of mgrB can result in oscillations in the two component system. A - Model schematic of an *in silico* mutant expressing *mgrB* constitutively instead of being expressed from the PhoP-P dependent promoter P_mgrB_ B - Simulations of PhoP-P following a switch from high (50 mM) to intermediate (1 mM) Mg^2+^show oscillations if *mgrB* is expressed at constant but low levels. Oscillations are absent in wild-type models, as well as models expressing *mgrB* at a constant high rate.

To understand the mechanism of the oscillations and why elimination of MgrB feedback create these we constructed a simplified model of the circuit (see S3 Text for detailed analysis). The result showed that oscillations arise from autoregulated expression of *phoPQ* operon, namely from PhoP-P dependent increase in PhoQ concentration. Indeed, since PhoQ is a bifunctional enzyme, autoregulation results in simultaneous positive and negative feedback [24,25]. However, if PhoQ is first produced in a kinase conformation [16] and slowly switched to a phosphatase conformation, there will be a time-delay between the positive and negative component of the feedback. The fast positive and slow negative feedback leads to oscillations. Increase in MgrB concentration increases the fraction of PhoQ that is MgrB bound, speeding up PhoQ conversion to phosphatase state and thereby prevents oscillations by reducing the delay in the negative feedback. This is why oscillations are not observed when MgrB is upregulated by PhoP-P or when constitutive production of MgrB is too high (Fig 6B). Thus, we find that autoregulated PhoPQ TCS may use negative feedback through MgrB in order to avoid sustained oscillations in response to stimulus. Notably the oscillations are not a consequence of any particular model assumption, but rather seem to stem from a general mechanism that can be applicable to many autoregulated TCSs. It remains to be seen if this mechanism can lead to oscillatory or pulsatile response for systems where it is physiologically beneficial.

## Discussion

The output of some two-component systems with bifunctional kinases – phosphorylated response regulator protein – displays robustness to overexpression of the two proteins. Here we show that this property can extend to PhoPQ TCS which regulates the gene encoding MgrB that inhibits PhoQ kinase activity resulting in a negative feedback loop. Using models of PhoPQ TCS we show that PhoP-P can be insensitive to overexpression of PhoPQ if MgrB is expressed in excess of PhoQ. The PhoPQ TCS steady state response to decreasing Mg^2+^concentration displays two distinct phases of activation separated by a plateau. We propose roles for Mg^2+^and MgrB in modulating PhoQ activity such that a model recreates the biphasic nature of steady state dose-response. We propose that Mg^2+^binds to PhoQ and promotes the phosphatase conformation. Limitation of Mg^2+^then drives a change in conformation of PhoQ to the kinase form. We also hypothesize that MgrB suppresses the rate of this conformation change and the autokinase activity of PhoQ. Next, we find approximate analytical solutions for PhoP-P at different ranges of Mg^2+^concentration. In our models we find that strong MgrB-mediated suppression of rate constants and growth-dilution of proteins are important factors that shape the biphasic dose-response. Finally, we propose advantages gained by having such an overlaid feedback structure. Negative feedback limits activation at low but not growth limiting Mg^2+^and shapes a surge in transcription in response to large step downs in Mg^2+^. Whereas positive feedback enables a strong activation of PhoP-P dependent promoters at growth-limiting Mg^2+^.

With a third component modulating the kinase’s activity, how does a PhoPQ TCS still obtain robustness to levels of the two components? Our analysis shows a possible condition in which PhoP-P can be robust to variations in PhoPQ expression. If MgrB is expressed at much higher levels compared to PhoQ, dependence of PhoP-P on total PhoPQ expression becomes negligible. This condition for robustness is not implausible within *E. coli* since *P_mgrB_* is one of the strongest PhoP-activated promoters [16] and is likely weaker than *P_phopQ_*. PhoQ, like many TCS sensor kinases, is expressed at low concentrations, estimated 50 fold less than PhoP [19,23]. In fact, measurement of protein synthesis rates puts the ratio of MgrB to PhoQ between 3 and 23 depending on the culture medium [23]. It is possible that over the range of induction rates (upto 4x wild type) MgrB remains in excess of PhoQ.

Multiple PhoP-P dependent promoters plateau over a range of 1 to 0.01 mM Mg^2+^while remaining far from saturation [9], only to be stimulated strongly when Mg^2+^becomes growth limiting (< 10^−3^*mM*). How does PhoPQ output plateau at lower stimulus levels, but still maintain the ability to respond strongly when needed? Hypotheses of Mg^2+^binding PhoQ at multiple sites with differing affinities have been made [9], however our analysis uncovers a potential mechanism with fewer assumptions. Consistent with our assumptions, studies in *Salmonella* strains have suggested that Mg^2+^binding PhoQ increases its phosphatase activity. We propose a model in which Mg^2+^directly binds a kinase-conformation of PhoQ and switches it to phosphatase-conformation. Our analysis shows that if MgrB strongly suppresses autophosphorylation, as well as the switch from phosphatase to kinase conformation, dilution due to growth remains the only way through which PhoQ-MgrB phosphatase complex can decrease. This can create a regime where both phosphorylation and dephosphorylation of PhoP are independent of the signal rate (switching from kinase to phosphatase).

Finally, what are the advantages of encoding a negative feedback to limit activation of an autoregulated two-component system? PhoPQ two-component system is widely conserved across bacterial species including pathogenic bacteria such as *Yersinia pestis* and *Salmonella typhimurium* [10,14,18]. Interestingly, PhoP-P regulons in these species also encode molecules that limit PhoQ activity [18]. Conserved structure of the network suggests that the structure provides some fitness advantages in Mg^2+^limitation by controlling level of activation of PhoP-P regulon. In *E. coli* positive autoregulation of the PhoPQ TCS helps cells survive in growth-limiting Mg^2+^. On the other hand, negative feedback creates a transcription surge in response to a step down in Mg^2+^concentration. Negative feedback also facilitates a faster response compared to a mutant strain expressing *mgrB* constitutively at levels such that steady state response of the two strains is comparable [16]. Other negative feedback designs can also provide some of the same benefits. Phosphate sensing PhoBR TCS in *E. coli* speeds response by encoding a negative feedback. In contrast with PhoPQ, PhoB-P does not upregulate a protein that suppresses kinase activity. Instead, PhoB-P represses the autoregulated *phoBR* promoter at high concentrations of PhoB-P [26]. This design can overcome the costs of positive autoregulation; however, this design does not create a transcription surge.

In *Salmonella*, benefits of a surge in transcription can be obtained independent of negative feedback through an MgrB-like protein post-translationally suppressing kinase activity [11,24]. Models of the *Salmonella* PhoPQ TCS reveal how positive autoregulation and the phosphatase activity of PhoQ together create an initial surge and a later decrease in expression of genes in PhoP-P regulon. Interestingly, models are consistent with the observed loss of surge in transcription if *phoPQ* is expressed constitutively. In contrast, transcription surge is present with either constitutive or autoregulated *phoPQ* expression in *E. coli* [16]. The surge and subsequent decrease in transcription is lost in strains lacking *mgrB*, suggesting different mechanisms drive transcription surges in *E. coli* and *Salmonella*.

Post-translational negative feedback also helps maintain sensitivity of PhoQ over a wider range of stimulus levels. The *E. coli* PhoBR TCS does not show adaptation in response to a downshift in phosphate. The advantage of overlaid autorepression and positive autoregulation in PhoBR TCS is that it allows for selecting a stronger autoregulated promoter without sacrificing speed [26, 27], but limits maximum activation. For two TCSs that sense low level of two nutrients, what selection pressure could have led to evolution of these structurally similar but functionally different negative feedback loops remains unknown. Taken together, these findings show how interplay of positive and negative feedback can shape dynamical properties of the PhoPQ two-component system.

Using our models of PhoPQ TCS that can explain experimental observations, we can make some testable predictions. First, if the experiment measuring TCS output as a function of independent induction of *phoPQ* operon (Fig 1B) is conducted using a strain expressing *mgrB* constitutively, we predict that robustness should not be observed (Figure 2 in S3 Text). This lack of robustness of PhoP-P output to PhoPQ overexpression could become more apparent if *mgrB* is expressed at low levels. Second, with the same strain expressing *mgrB* constitutively, we find that oscillations are possible in the output when cells are switched from high to intermediate Mg^2+^, eg. 50 → 1mM (Figure 1 in S3 Text). Oscillations are not predicted if the cells are switched to low Mg^2+^, i.e. 0.01mM or if *mgrB* is expressed at high levels (comparable to expression levels of wild-type cells at 0.01 mM Mg^2+^, S4 Fig). Moreover, we observe these oscillations with *in silico* mutants of wild-type models that show a biphasic dose-response. The predictions of oscillation are relatively robust and no oscillations are observed with models that fit temporal data well but fail to display biphasic dose-response. These predictions must be tested in the future.

Notably, the uncovered oscillation mechanism is quite generic. It requires transcriptionally autoregulated TCSs with sensory kinase in two conformations, one kinase-dominant another phosphatase dominant. If both kinase and phosphatase states of the sensor kinase (SK) are increased proportionally with the increase in total-SK, the steady state RR-P is independent of total-SK, as has been seen in multiple experimental and theoretical works [20–22]. In other words, positive feedback (increase in kinase form of SK) exactly balances out a negative feedback (increase in phosphatase form). While the above argument may hold true about steady state RR-P, positive and negative feedback may have different timescales. If sensory kinase in produced in the kinase-dominant conformation and then slowly switches to phosphatase-dominant one, sustained or damped oscillations are possible. Most natural TCSs are autoregulated and often activated by a ligand that drives a conformational change in sensory kinase [28,29]. Therefore this oscillatory or pulsatile response dynamics may be observed in the systems where it is of physiological benefit and could be used in synthetic biology applications.

## Materials and methods

### Model and simulations

Two mathematical models were developed to examine the dynamical properties of PhoPQ TCS. The first model considers a single bifunctional form of the kinase PhoQ, whereas the second model considers two separate conformations (kinase and phosphatase). A set of ordinary differential equations (ODEs) describes the rate of change of all protein and mRNA species (S1 Appendix). The phosphorylation and dephosphorylation cycle reactions follow previous models by Goulian and collaborators [19]. Gene transcription regulation is modeled by phenomenological models of (Hill-function) dependence on [PhoP-P].

All models used (S1 Appendix) were simulated to follow experimental protocol as closely as possible. For time course, signal parameter (depending on model) was set to 1mM to compute an initial steady state vector of all state variables using ode15s in MATLAB. Using this as initial condition, signal was set at a pre stress value (50mM or 2mM) and integrated for 3.5 hrs. Then the signal was set to a post-stress value (0.01mM, 2mM or 10mM) and integrated for 2 hrs, at the same time points as the data. For steady state data, signal was set to the respective value and integrated to steady state.

### Experimental data

Experimental data was obtained from refs [16, 19]. Time course of YFP:CFP read out from plates following a switch from 50mM to 0.01mM Mg^2+^published in ref [16] was used to fine tune temporal parameters. YFP was either expressed from the PhoP-P dependent *mgrB* promoter or *phoPQ* promoter. Time course data for the following strains was used as constraint on parameters: wild-type (*P_mgrB_, P_phoPQ_*), *mgrB* deletion (*P_mgrB_, P_phoPQ_*), autoregulation deletion, autoregulation+ *mgrB* double deletion. In addition, wild-type time-course data collected from *P_mgrB_* promoter for cells switched from 50mM to 10mM, 50mM to 2 mM and 2mM to 0.01 mM was also used. The value of YFP:CFP at t = 0 for wild type cells switched from 50mM to 0.01mM with YFP expressed from *mgrB* promoter was used to normalize all time-course data. Steady state YFP:CFP data with YFP expressed from *P_mgrB_* for a range of Mg^2+^concentrations (30mM to 0.03mM) published in ref [19] was used to fine tune steady state parameters. This steady state data was normalized to YFP:CFP at 30mM. All experimental data used for fitting is shown in S2 Fig.

### Error calculation

#### Time course

If (*t_i_, y_i_*) is the experimental normalized YFP:CFP data for a given strain and (*t_i_, y_i_*) represents simulated normalized YFP:CFP, then the squared residual error for time course is calculated as

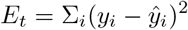

Similar residual errors for all strains are then added together to give a total time course residual error *E_t_*.

#### Steady state signal response

Steady state simulated YFP:CFP at each signal level is normalized to the simulated value at 30mM 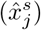 and compared against the experimental value 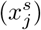 leading to the residual error

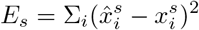

In addition, steady state YFP:CFP is simulated at very high signal (equivalent of 10^−4^mM). If this value is not > 6× [YFP:CFP(0.03mM)], then an error penalty of 25 (comparable to the maximum squared residual at 0.03 mM) is added to *E_s_*.

### Parameter fitting

Parameters were fit to minimize the above squared residual error (*E_s_ + E_t_*) using particle swarm optimization in MATLAB. Each particle swarm optimization run resulted in one parameter set. The best fitting parameter sets were used for further analysis. All codes and parameter/data sets can be found in the following GitHub repository: https://github.com/satyajitdrao/PhoPQManuscript.git

## Supporting information

**S1 Fig.**
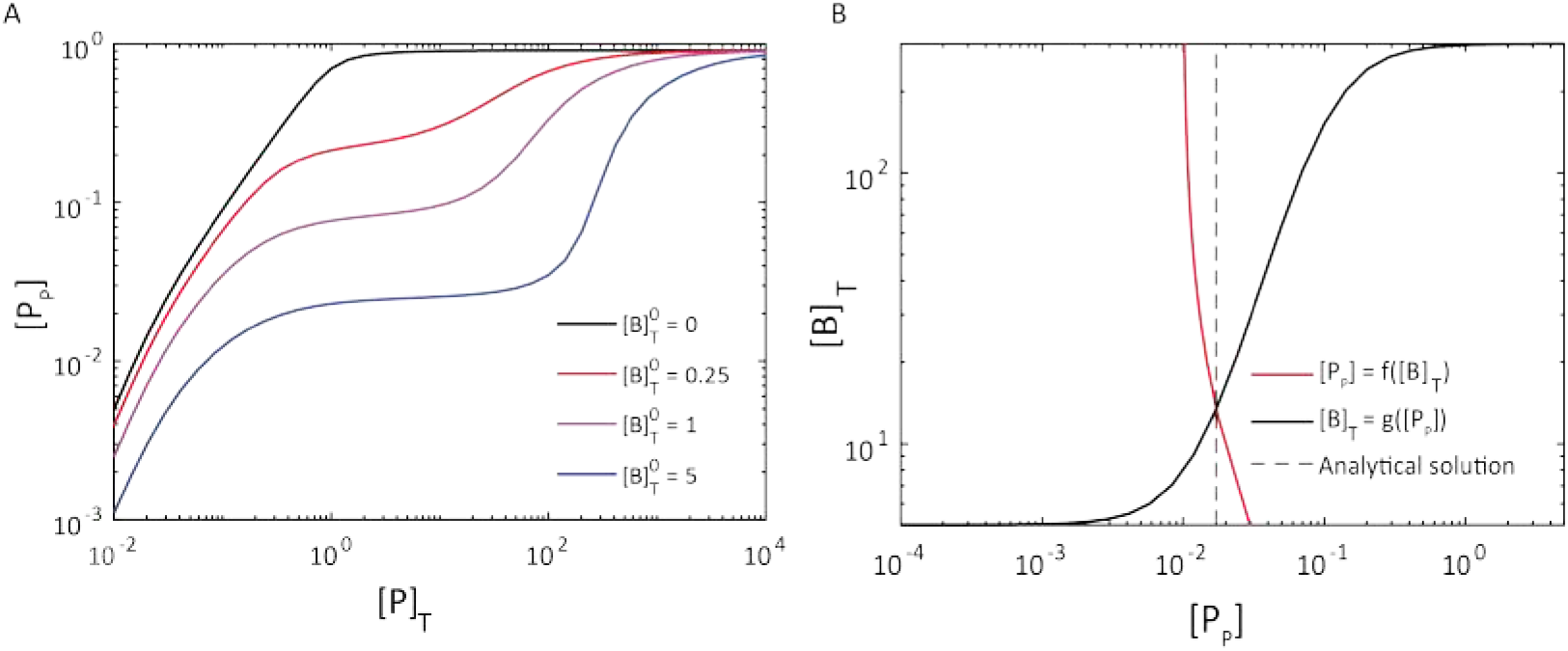
Steady state simulations of one-state PhoQ model of PhoPQ TCS. Steady state [PhoP-P] as a function of total PhoP, PhoQ at various 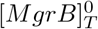 levels (all concentrations in units of (*C_P_ + C_t_*). As 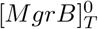 increases, a range of total-PhoP,PhoQ appears where PhoP-P is robust (B) Each point in (A) is an intersection of transcription and interaction modules (Figure 2A, main text). Red curve shows solution to the interaction module. At fixed total PhoP,PhoQ, total-MgrB is increased and steady state [PhoP-P] is computed. Black curve represents the saturating dependence of MgrB-total on [PhoP-P]. The dotted line is the analytical solution obtained by solving equations 1 and 2 in main text

**S2 Fig.**
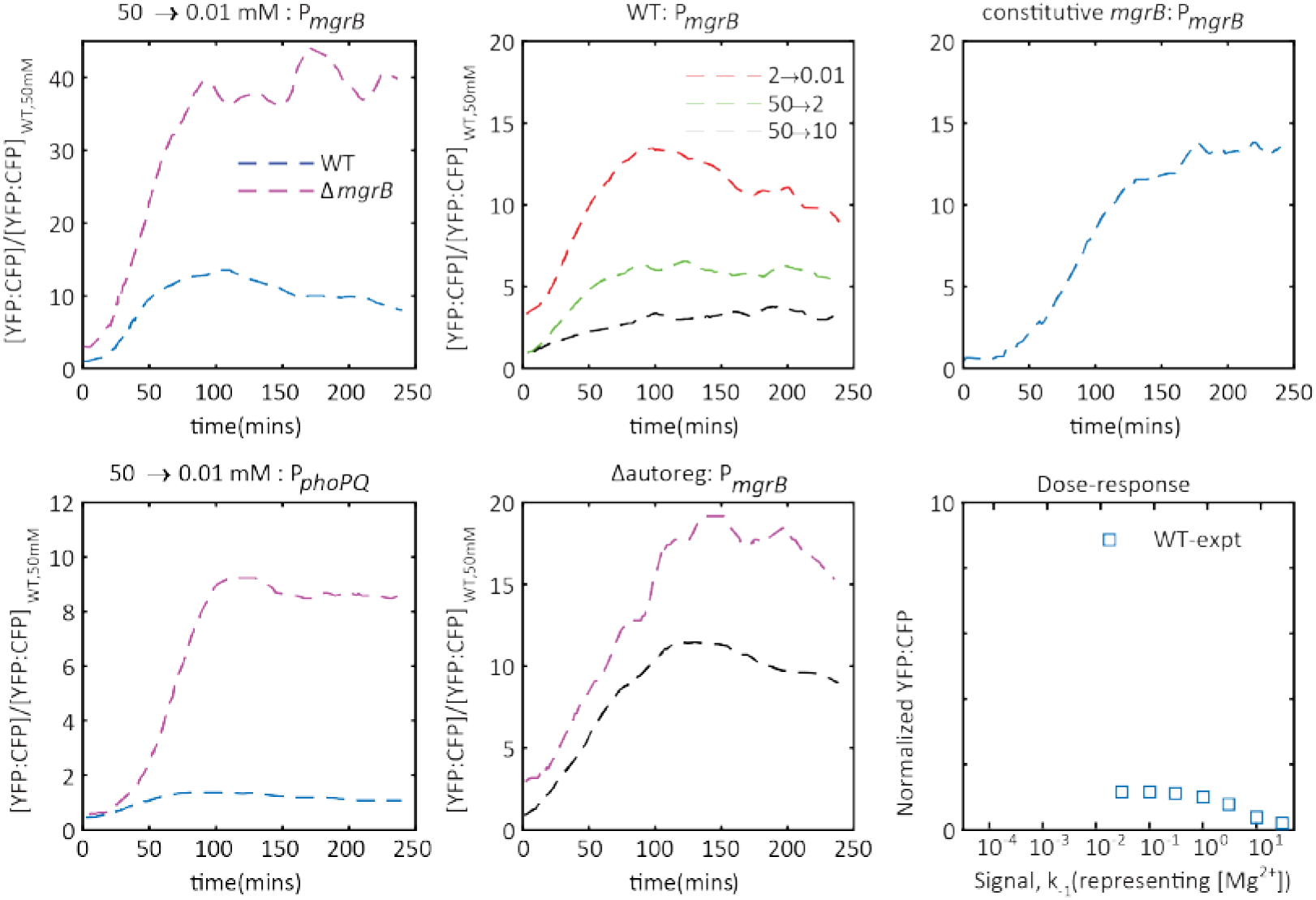
Experimental data used for parameter fitting. YFP:CFP ratio (normalized to the ratio at 50mM) extracted from [16] and [19]. This normalized data set was used to fit temporal and steady state parameters for all models described in this paper. Data was extracted using image analysis in MATLAB (except figure f, which was read out manually from ref [19]). All data except constitutive *mgrB* (top right panel) was used to fit models

**S3 Fig.**
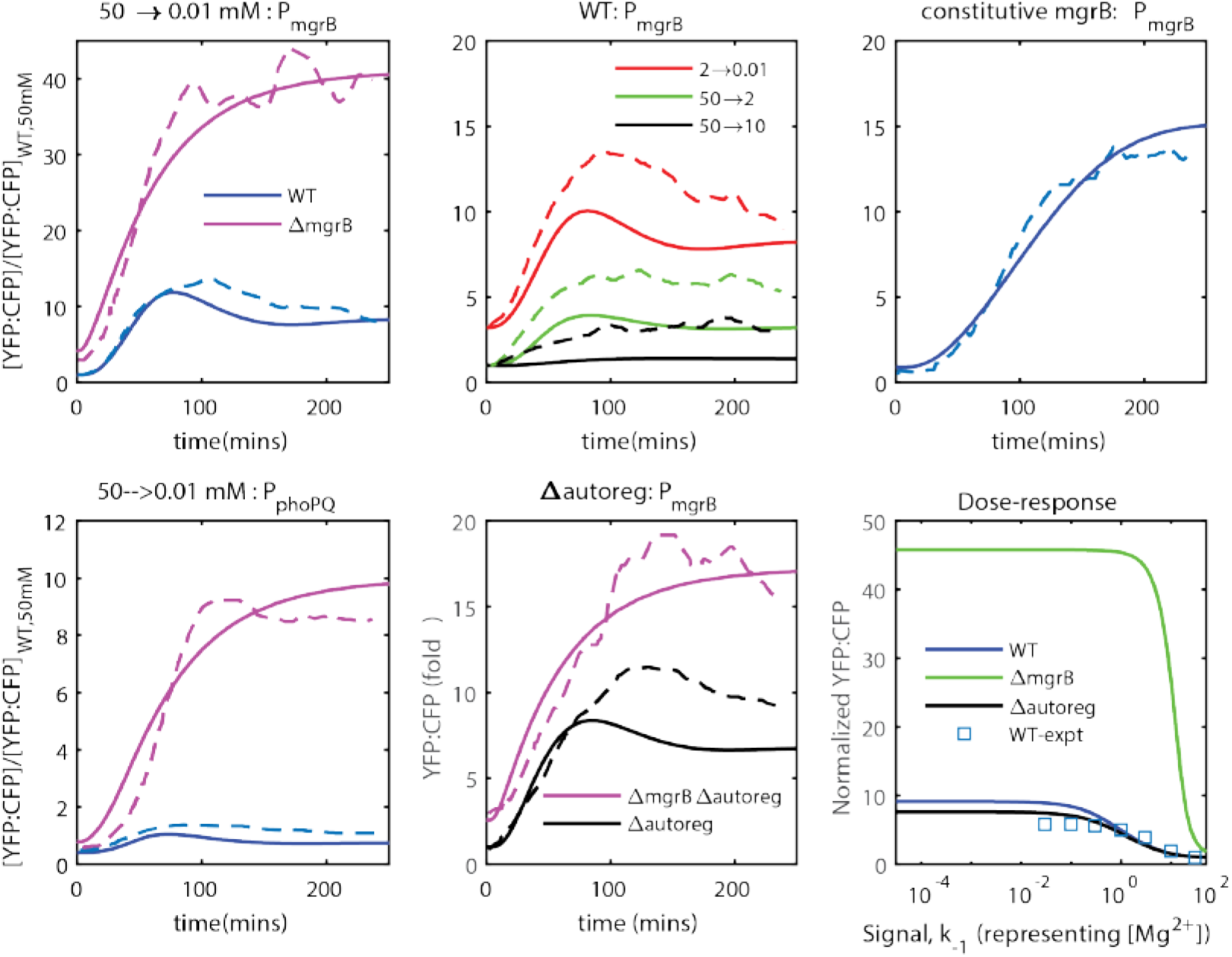
Best fit simulations of dynamical model 1. Simulations from a representative parameter set showing best quantitative fit for the simple PhoPQ model. Simulation of an in-silico mutant expressing *mgrB* constitutively (top right, solid line) at 10x basal transcription rate of *mgrB* in wild-type. This simulation in addition to PhoQ phosphatase-lacking mutant was used to verify whether a parameter set was accurate

**S4 Fig.**
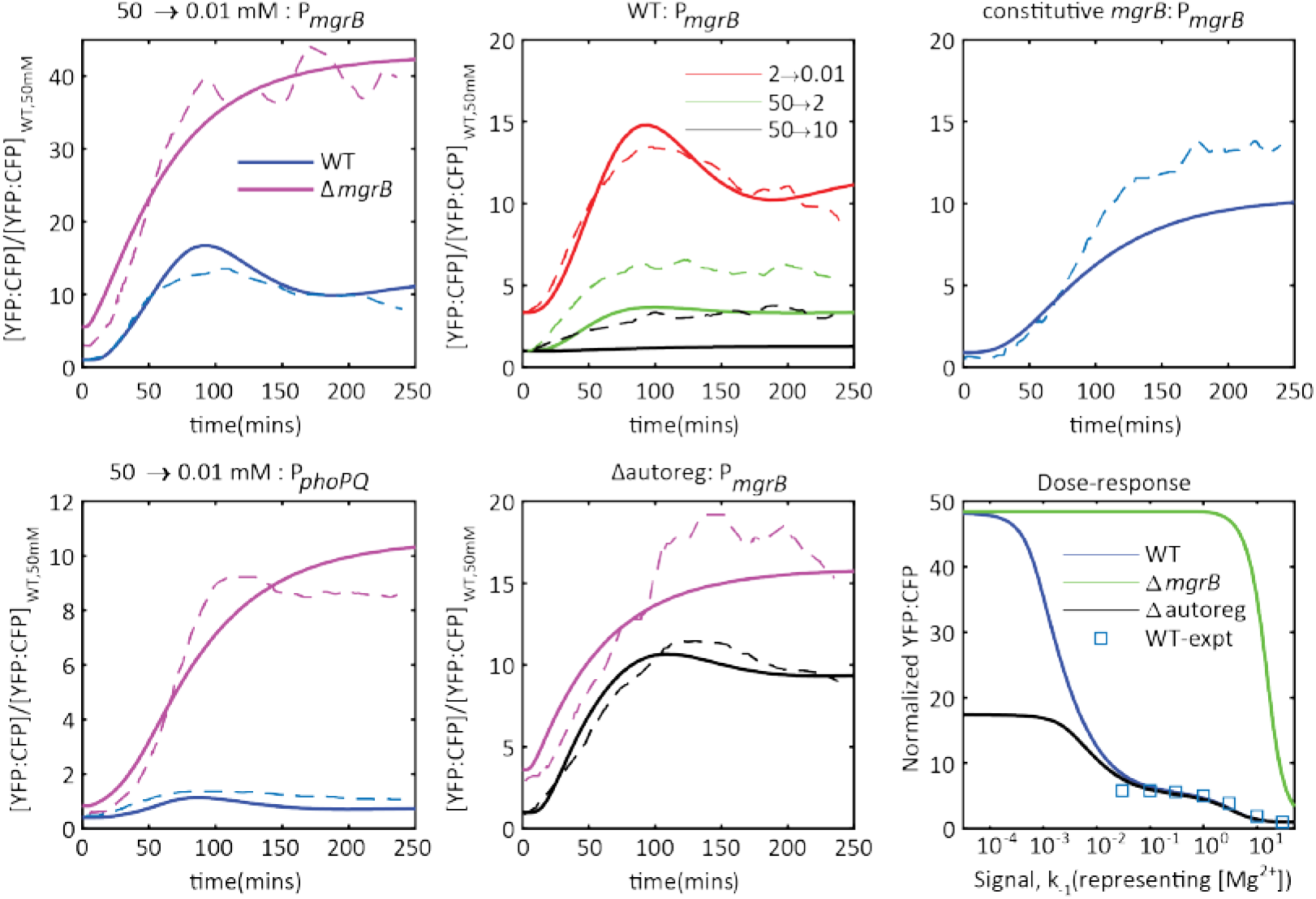
Best fit simulations of dynamical model 2. Simulations from a representative parameter set showing best quantitative fit for the two-state PhoPQ model

**S5 Fig.**
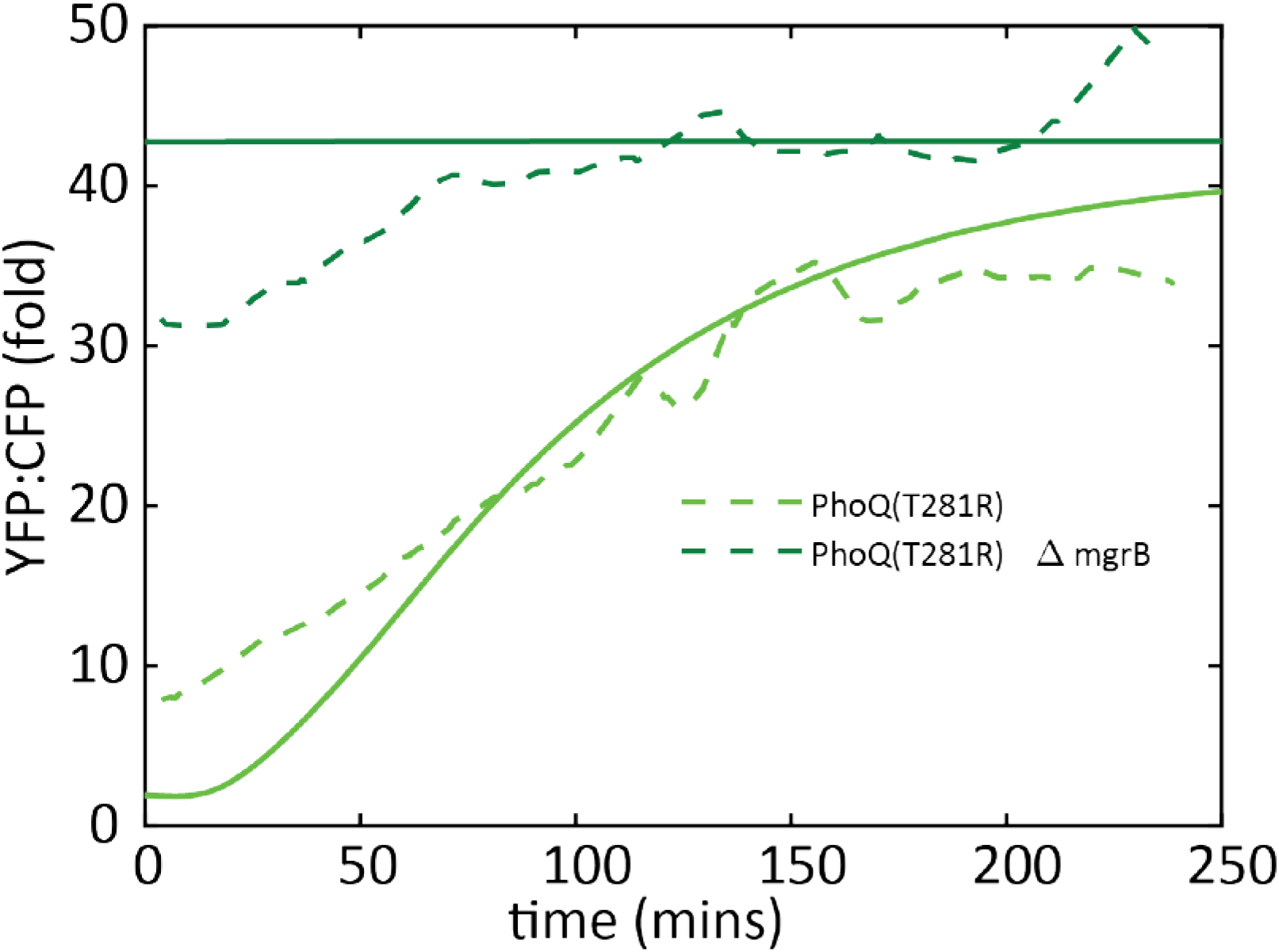
Verifying accuracy of dynamical model 2 by simulating phosphatase lacking *in-silico* mutant of PhoQ. Verifying accuracy of parameters by simulating PhoPQ TCS model without PhoQ phosphatase activity. With parameters that generate the fit in S4 Fig, we simulate the Mg^2+^step-down with an *in-silico* mutant of PhoQ lacking phosphatase activity single mutant (light green, solid line) or double mutant with *mgrB*-deletion (dark green, solid line). These simulations show a qualitative match with the experimental data (dashed lines).

**S1 Text. Analyzing PhoPQ-MgrB reaction network for absolute concentration robustness**

**S2 Text. Steady state PhoP-P for one-state PhoPQ-MgrB model**

**S3 Text. Model predictions and analysis of oscillations**

**S4 Text. A framework to examine steady state signal response for two-state model of PhoPQ-MgrB**

**S1 Appendix PhoPQ TCS models: Reactions, ODEs and Parameters**

## S1 Text: Analyzing PhoPQ-MgrB reaction network for absolute concentration robustness

Biochemical reaction networks can possess a property of absolute concentration robustness (ACR) based on the structure of the network alone [1]. If a biochemcial reaction network satisfies certain conditions related to the structure of the network, ACR can result in the concentration of one species being robust to variation in all other species concentrations. Here we analyze whether the interactions in PhoPQ TCS satisfy the conditions for ACR. PhoQ and all its forms are represented by Q and MgrB is represented by B for readability and ease of writing. The biochemical reaction networks shown in Figure 1 are analyzed for properties such as nodes, strong linkage classes and rank. The theorem given in ref [1] was used to identify absolute concentration robustness.

**Figure 1:**
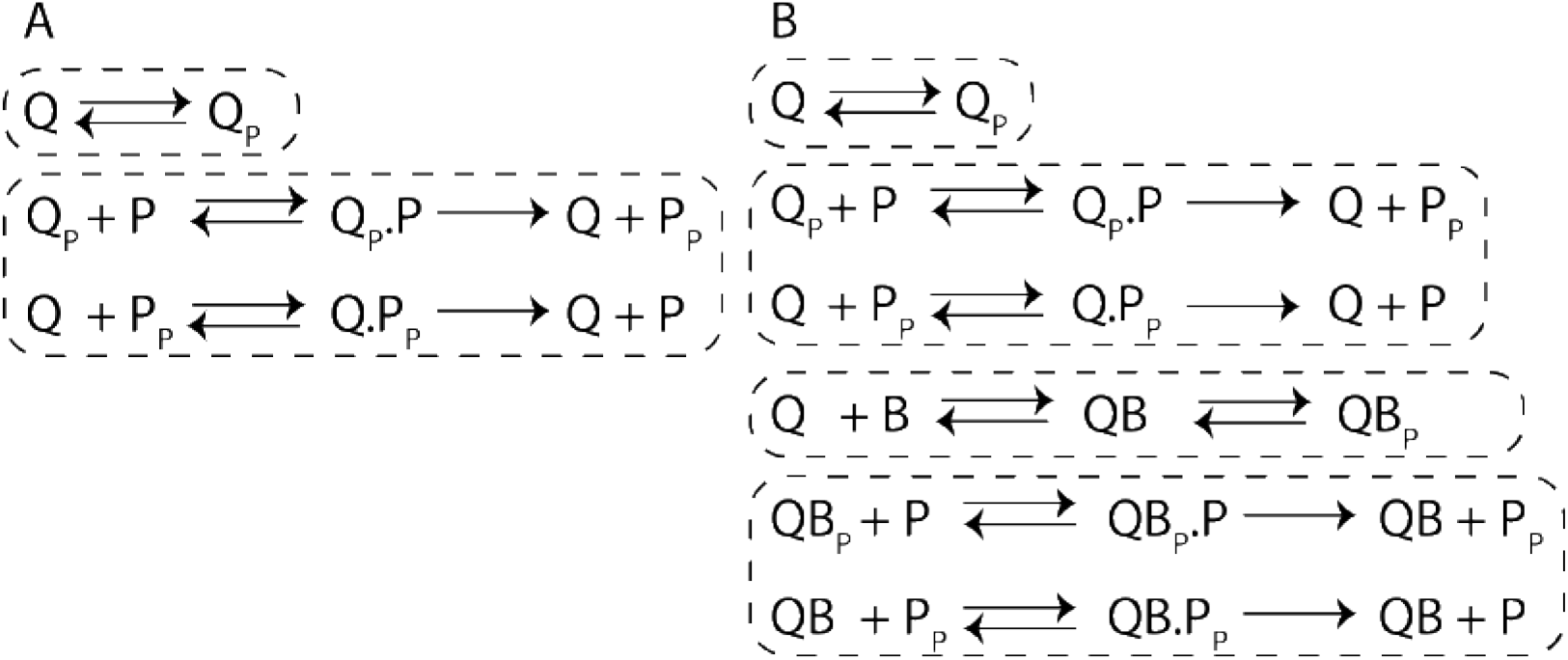
Dashed rectangles indicate strong linkage classes. Biochemical reaction network for a typical TCS (A) compared to a TCS with a third component B binding Q and affecting subsequent catalytic activity (B)

**Table 1:**
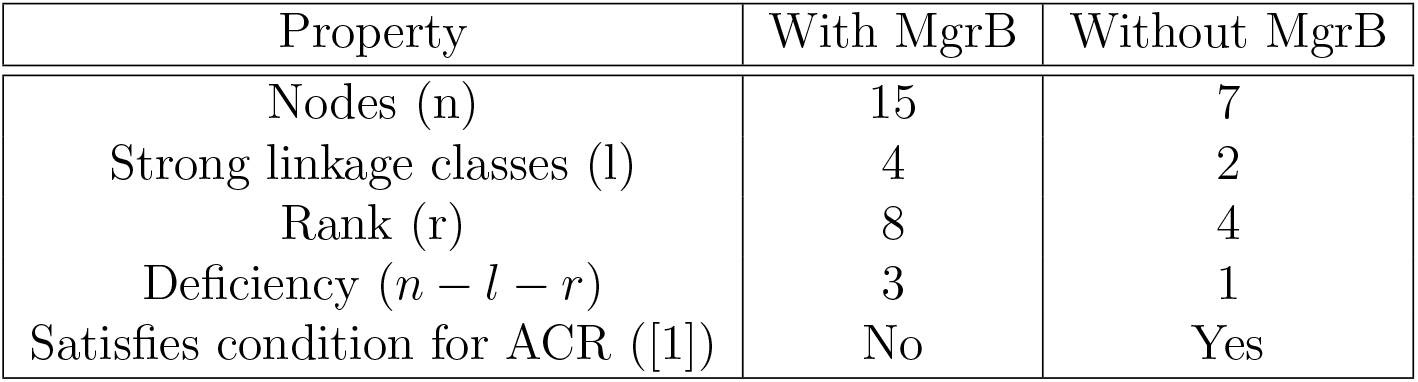
Network-analysis of PhoPQ TCS interactions

| Property | With MgrB | Without MgrB |
| --- | --- | --- |
| Nodes (n) | 15 | 7 |
| Strong linkage classes (l) | 4 | 2 |
| Rank (r) | 8 | 4 |
| Deficiency ($n - l - r$) | 3 | 1 |
| Satisfies condition for ACR ([1]) | No | Yes |

## S2 Text: Steady state PhoP-P for one-state PhoPQ-MgrB model

In this section we identify conditions under which PhoP-P is robust to over-expression of PhoP and PhoQ (Figure 2, Main text). We find an approximate analytical solution to This model is the simplest modification of the PhoPQ model created by Miyashiro and Goulian in [3]. Reactions and ODEs are described in Supplemental File 4. Phosphotransfer and phosphatase complexes at steady state give:

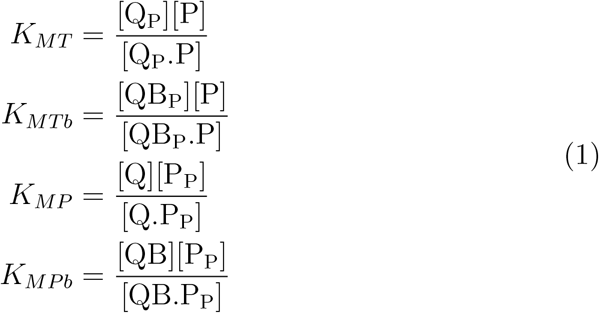

*K_MP_, K_MPb_, K_MT_, K_MTb_* are Michaelis-Menten constants for the catalytical reactions shown in Figure 1 S1 Appendix. At steady state, the flux of phosphorylation of PhoP can be matched with the flux of dephosphorylation of PhoP-P. This gives

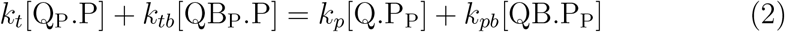

Substituting complex concentrations from Equation 1:

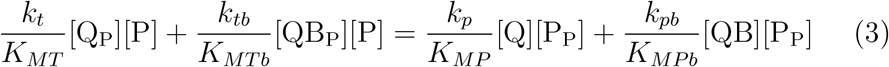

Steady state expressions for [Q-P] and [QB-P] can be found by first setting 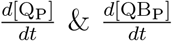 to zero, followed by plugging in complex concentrations from 1. The expressions can then be plugged into Equation 2.

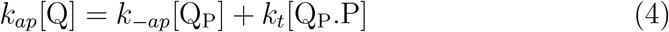

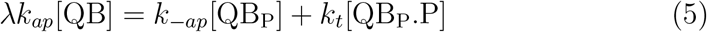

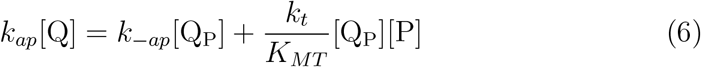

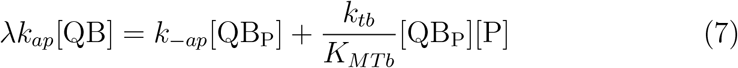

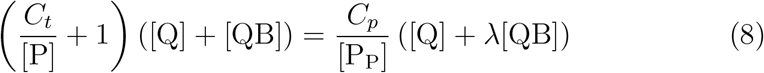

The expression for [QB] in terms of [Q] and [B] can be found by setting 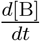 to zero.

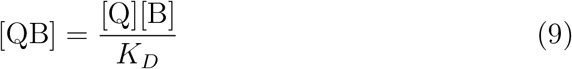

Plugging back into Equation 8,

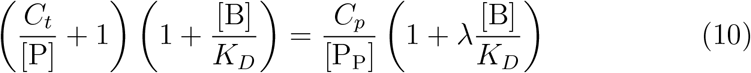

Where 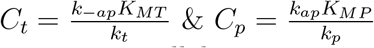.

This equation will be constrained by conservation equations for PhoP, PhoQ and MgrB. In general,

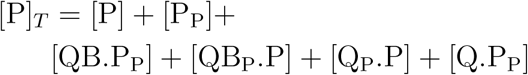

However, two-component system histidine kinases are usually expressed at much lower concentrations compared to response regulators. Ratio of PhoP:PhoQ in *E. coli* is reported to be much larger than 1 [2, 3]. Therefore:

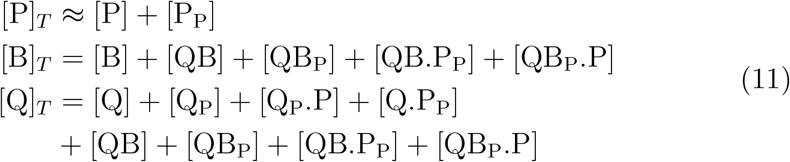

Negative transcriptional feedback dictates the following saturating relation between B-total and PhoP-P

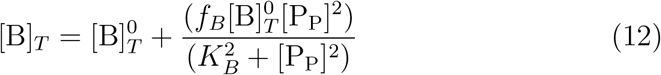

The set of equations 10, 11, 12 together form the complete model of PhoPQ TCS. If we set MgrB-total to zero, this set of equations reduces to the model in ref [1] and the concentration of [PhoP-P] is the negative solution of a quadratic as discussed in refs [1,3,5]. In general however, we solve equations 10, 11, 12 for [PhoP-P] assuming the following parameters: *C_p_* = 10*C_t_, K_D_* = 0.1, λ = 0.01, *K_B_* = 5. *f_B_* was measured in ref [4] at ~60. 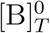 was varied from 0 to 5 (S1 Fig). All concentration variables in units of (*C_p_ + C_t_*). At each 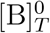, the equations were solved for a range of P-total values ([Q]_*T*_ = [P]_*T*_/50) as shown in S1 Fig. Non-linear equations solved using fmincon function in MATLAB to obtain steady state PhoP-P.

At high levels of [B]T, we can rewrite total MgrB conservation in equation 11 as [B]_*T*_ = [B] + *ϵ*, where e represents terms of the order [Q]_*T*_/[B]_*T*_. At the limit *ϵ* → 0, [B] ≈ [B]_*T*_. Equations 11 can then be incorporated in a straightforward way into eq 10.

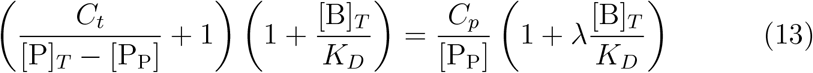

As [P]_*T*_ increases, the term *C_t_*/[P] (dependent on autodephosphorylation) becomes less significant as explained previously in ref [1], leading to robustness of [PhoP-P] to total-PhoP,PhoQ. In the special case that *C_t_* = 0 which happens at *k*_−ap_ = 0, [PhoP-P] outright does not depend on total-PhoP as discussed in detail in ref [5].

## S3 Text: Model predictions and analysis of oscillations

**Figure 1:**
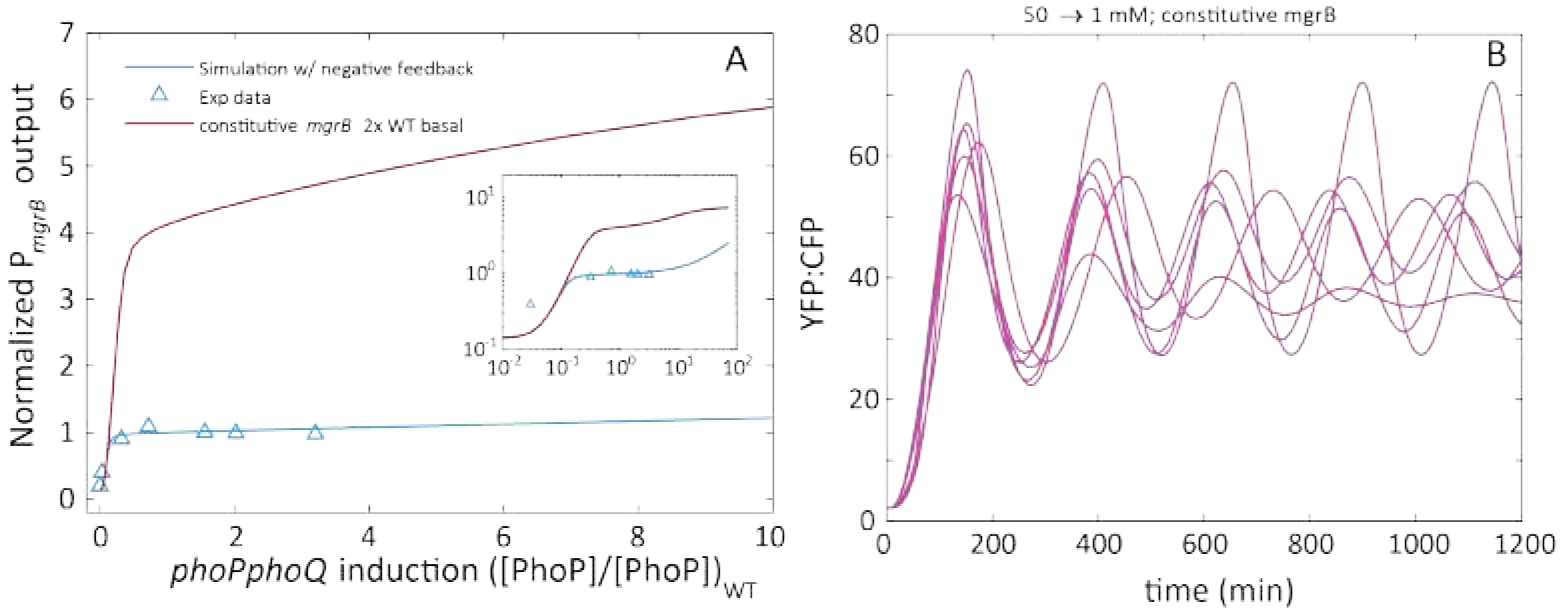
A - PhoPQ TCS models predict that PhoP-P output will not be robust to overexpression of PhoP and PhoQ if *mgrB* is expressed constitutively. Blue triangles and blue line represents experimental data and simulation of PhoP-P output to overexpression of *phoPQ* operon as described in [1]. Red line shows PhoP-P output as a function of *phoPQ* induction if *mgrB* is expressed from a constitutive promoter at twice the rate of the wild-type (at basal activity level). Inset: Models with *mgrB* expressed from *P_mgrB_* will show an increase in PhoP-P output at 100 fold overexpression of *phoPQ* B. Following a switch in Mg^2+^concentration from 50 to 1 mM, our PhoPQ models with constitutively expressed *mgrB* (at levels comparable to wild-type basal) predict oscillations in PhoP-P output. Each line represents a parameter set that explains wild-type data (including biphasic dose-response) as shown in S4 Fig

In order to understand the role of negative feedback in shaping the dynamical properties of the TCS, we constructed *in-silico* mutants of our wildtype models with negative feedback through *mgrB*-upregulation removed. We constructed *in-silico* mutants expressing *mgrB* constitutively. We find that models of wild-type PhoPQ TCS that fit well with temporal data as well as display biphasic dose-response can show limit cycle oscillations in PhoP-P if *mgrB* is expressed at a constant rate comparable to the basal *mgrB* expression rates of corresponding wild-type models, i.e. if *mgrB* upregulation by PhoP-P is removed from the wild-type model. We observe limit cycle oscillations at signal levels equivalent to ~1 mM Mg^2+^(Figure 1B).

To understand the mechanism that results in limit cycle oscillations following removal of negative feedback through MgrB upregulation, we find a minimal model of the TCS that still retains oscillations (Figure 2). The minimal model consists of two conformations of the kinase (Qkin and Qph). Each conformation catalyzes phosphorylation or dephosphorylation reactions with a saturating dependence on the substrate, P and P_P_ respectively. We assume a constant amount of total-PhoP that is much greater than PhoQ as positive autoregulation of PhoP doesn’t contribute to the oscillations significantly. We find that positive feedback through upregulation of PhoQ (synthesized in a kinase-biased conformation Qkin; Figure 2), together with a slow negative feedback through conversion of Qkin to the phosphatase conformation (Qph) is sufficient to yield oscillations. In the wild-type model, we find that upregulation of MgrB by PhoP-P reduces the delay by speeding up kinase to phosphatase switch, as well as increases the total amount of MgrB bound PhoQ (which contributes largely to phosphatase activity rather than kinase activity).

**Figure 2:**
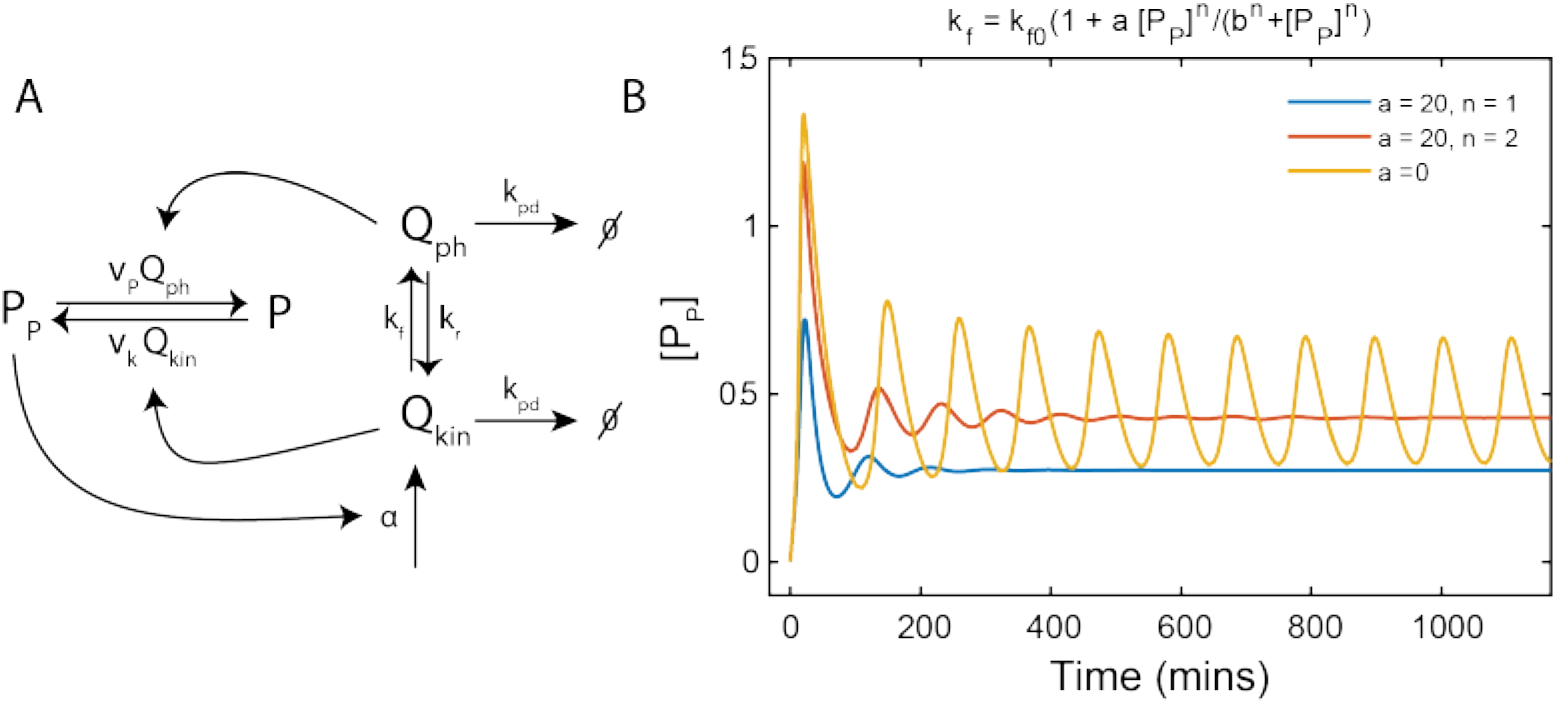
A - Minimal model of the model that yields oscillations in PhoP-P (P_*P*_). Phosphorylation and dephosphorylation reactions are modeled as Michaelis-Menten reactions catalyzed by Qkin and Qph respectively, i.e. 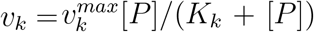, 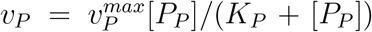. P_*P*_ drives synthesis of Qkin with a saturating dependence 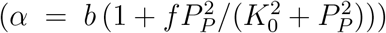. Qkin undergoes a reversible conversion to Qph. The rates used in this model were derived from the parameters used in the fully descriptive ODE model. B-The model yields limit cycle oscillations (blue). If the rate of kinase to phosphatase transition is modeled to increase as a function of P_*P*_ (to recapitulate effects of MgrB upregulation), the oscillations are suppressed and result in a stable steady state (red). Rates (units: *μM, s*): *k_pd_* = 3.1 × 10^−4^, *b* = 1.2 × 10^−6^, *f* = 26, *K*_0_ = 0.65, *v_k_* = 3, *K_k_* = 0.6, *v_P_* = 1.5, *K_P_* = 13.8, *P_T_* = 1.6, *k_f_* = 0.006, *k_r_* = 1.5 × 10^−6^

## S4 Text: A framework to examine steady state signal response for two-state model of PhoPQ-MgrB

In this section, we look at how this particular model (Fig. 3 Main Text, Figure 2 in S1 Appendix, S4 Fig) that fits well to experimental data is able to simulate a biphasic dose-response. Therefore, the analysis here is not general but dependent on the particular parameter set resulting in simulations that match experimental data (See Fig. 3 Main Text, S4 Fig). We make many assumptions along the way to attempt to find why PhoP-P can show a biphasic response to decreasing *k*_−1_. As we show in Fig. 4 (main text), the expressions we develop in this section for PhoP-P in ranges of *k*_−1_ corresponding to high (> 1 mM) and intermediate (1-0.01mM) Mg^2+^are good approximations of the steady state PhoP-P obtained from numerically solving the full model.

We analyze the model with no autoregulation, since that strain shows no difference in output compared to wild-type over the plateau region, both in experiments and in this model [1]. The mechanism for plateauing however remains valid even with autoregulation of *phoPQ*. Consider 4 forms of PhoQ: kinase, phosphatase and MgrB bound kinase, phosphatase. So the ODE system for PhoQ concentrations reduces to:

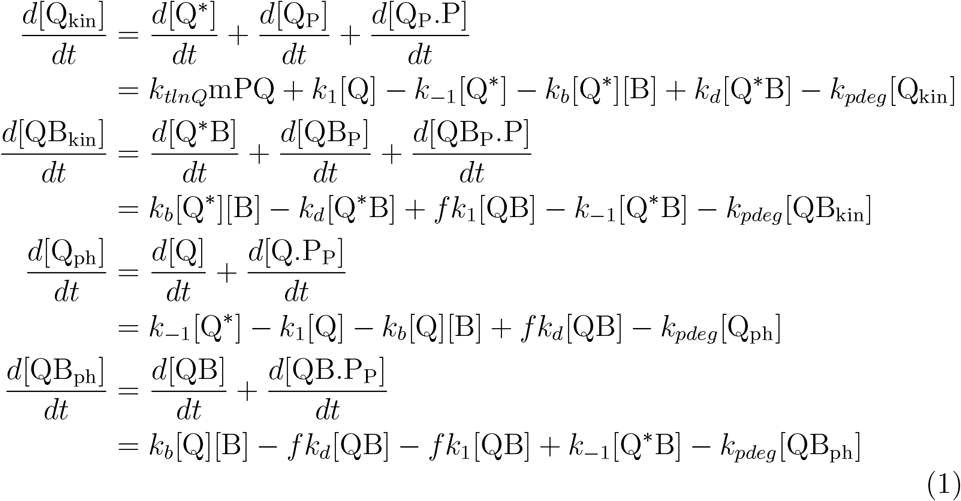

For simplicity, we do this analysis as λ →0 and f → 0. We assume all catalytic conversions within each form happen much faster than conversions between them. Thus the instantaneous concentrations of all sub-forms depend on the parent form in the following way:

- Kinase:

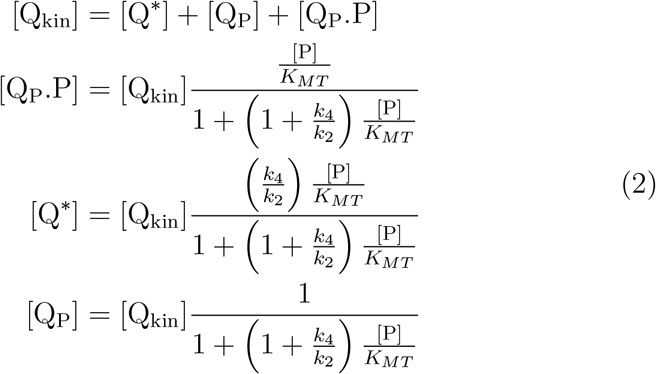
- Phosphatase:

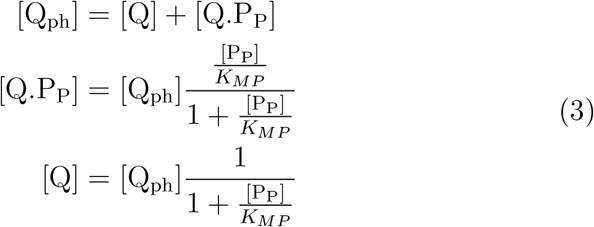
- MgrB-PhoQ Kinase:

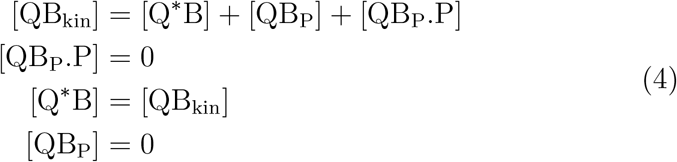
- MgrB-PhoQ Phosphatase:

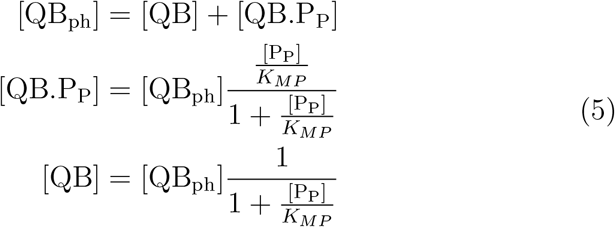

Now, we look at phosphorylation and dephosphorylation fluxes of PhoP to compute steady state PhoP-P.

- Dephosphorylation of PhoP-P

1. Phosphatase activity: Using paramters that generate optimal fit, we find that simulated [PhoP-P] remains well below the Michaelis Menten constant *K_MP_*. As a consequence, [Q_ph_] ≈ [Q] & [QB_ph_] ≈ [QB]. Moreover, at all signal levels contribution of Q_ph_ is negligible compared to QBph owing to most Q molecules being in bound (QB) state (Main text Fig. 4E).

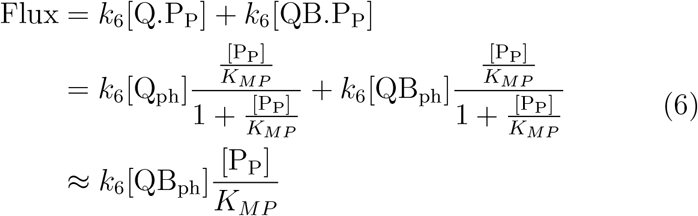
2. Flux due to growth-dilution = *k_pdeg_*[P_P_]
- Phosphorylation of PhoP: We assume flux of autophosphorylation is almost equal to phosphotransfer i.e. auto-dephosphorylation flux (*k*_−2_[Q_P_]) is negligible. Thus, PhoP-P formation flux ≈ *k*_2_[Q^*^]

At steady state,

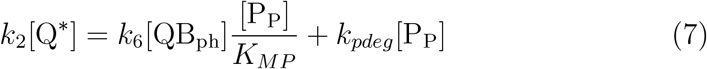

To compute [PhoP-P] at various signal levels (i.e. *k*_−1_ values) using this flux balance equation (Equation 7, we find approximate expressions for relation between [Q^*^] and [QB_*ph*_]. To this end, the set of equations in 1 can be solved at steady state to yield:

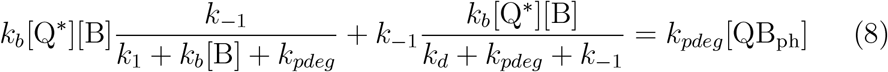

[B]_*T*_ = [B] + *ϵ*, where e represents total MgrB in PhoQ bound form. At all signal levels in our model, MgrB is in large excess of PhoQ, so that *ϵ* ~ 0. As a consequence, MgrB binding PhoQ or PhoQ* is a pseudo first order reaction with a rate of *k_b_*[B]_*T*_.

### High Mg^2+^

At high Mg^2+^ levels, PhoQ* deactivation (to PhoQ) is much faster than PhoQ* binding MgrB i.e. *k*_−1_ ≫ *k_b_*[B]. Whereas at both high and intermediate Mg^2+^levels deactivation rate is much larger than MgrB-PhoQ dissociation rate and growth-dilution rate i.e. *k*_−1_ ≫ *k_d_* + *k_pdeg_*. Moreover, rate constant for PhoQ* or PhoQ binding MgrB is much larger than activation rate of PhoQ to PhoQ* i.e. & *k_b_*[B] » (*k*_1_ + *k_pdeg_*). Therefore, the relation between [Q*] and [QB_ph_] i.e. Equation 8 reduces to:

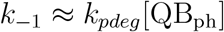

Plugging back into the flux balance equation i.e. Equation 7,

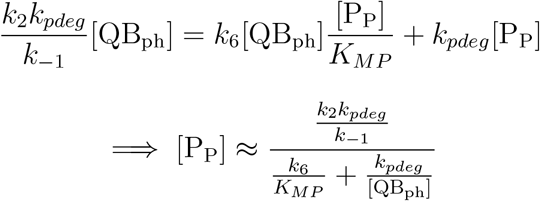

Since at this signal level [QB_ph_] [Q]_*T*_, and nearly insensitive to signal (Main text Fig. 4E), [PhoP-P] is inversely proportional to *k*_−1_, i.e. increases with signal. This equation can be plotted for values of *k*_−1_ corresponding to high range of Magnesium concentrations (Main text Fig. 4D, red dotted line).

### Intermediate Mg^2+^

At intermediate Mg^2+^levels, relation between the pseudo-first order rate constant of PhoQ* binding MgrB and PhoQ* deactivation rate constant is the opposite of high Mg^2+^i.e. *k*_−1_ ≪ *k_b_*[B]. Equation 8 can be approximated as:

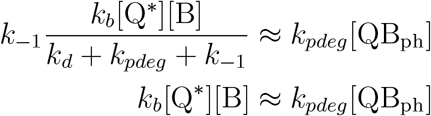

Since

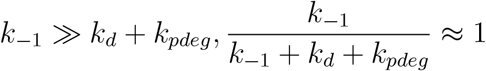

Plugging this relation between Q* and [QBph] back into the flux balance equation 7, we get

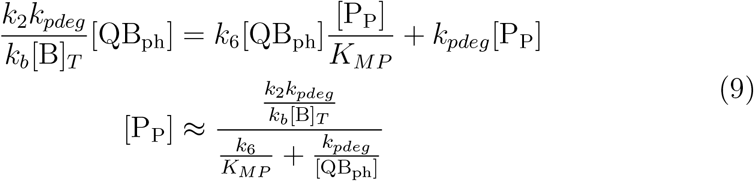

Over this range,[QB_ph_] can still be approximated as *Q_T_* and considered independent of signal (Main text Fig. 4E),

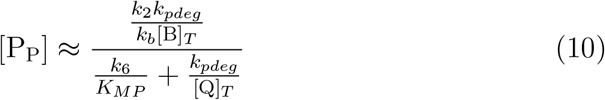

In this expression, there is no dependence on *k*_−1_, but on total MgrB. Total MgrB is a function of [PhoP-P], therefore steady state [PhoP-P] remains nearly independent of *k*_−1_(Main text Fig. 4D). Promoter output plateaus over this range (Main text Fig. 3). The plateau ceases to exist at lower Mg^2+^ as [QB_ph_] starts decreasing with signal (Main text Fig. 4E), and *k*_−1_ ~ *k_d_* + *k_pdeg_*. The equation above can be solved for a value of [P_P_]_*int*_ by solving the resulting cubic equation. Rearranging the above equation, [P_P_][B]_*T*_ = *c*, where 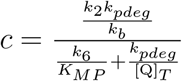. c is constant and independent of signal. [B]_*T*_ depends on [PhoP-P] as noted in Equation 12 S2 Text. This results in a cubic in [PhoP-P] which can be solved given the rate constants and value of [Q]_*T*_ to obtain [P_P_]_*int*_ (black dashed line, Main text Fig. 4D).

## S1 Appendix: PhoPQ TCS models: Reactions, ODEs and Parameters

### Model reactions: PhoPQ model with single bifunctional kinase

Shorthand - PhoQ = Q, MgrB = B

**Figure 1:**
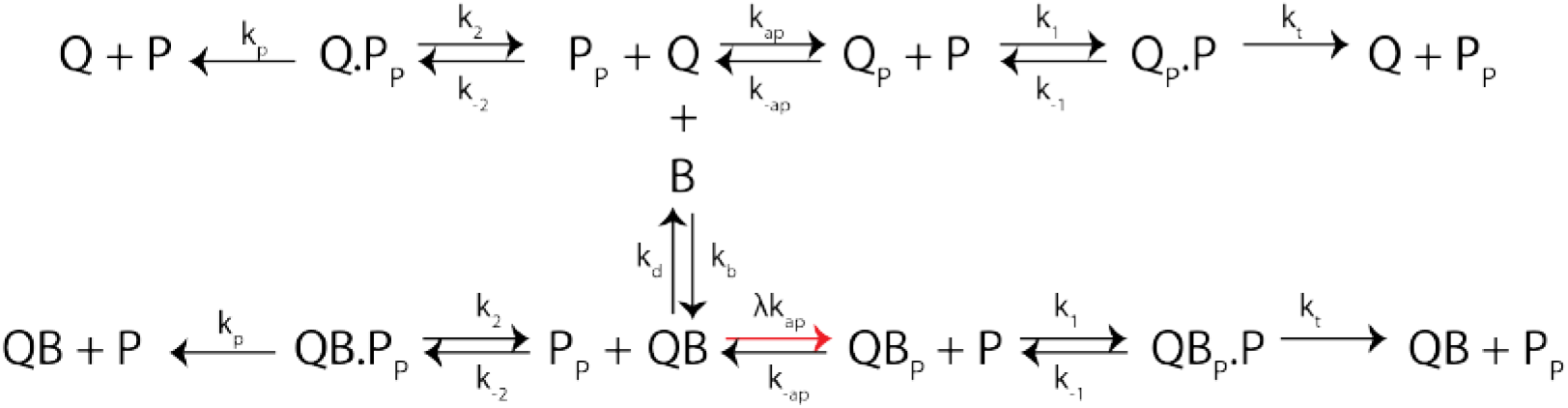
Figure shows all the reactions in PhoPQ TCS with associated rate constants. Reactions in the top row represent those in a typical TCS. This modified model includes Q binding B. QB undergoes all the same steps as Q (bottom row) with same rate constants, except autophosphorylation.

#### Post-translational reactions

- Signal sensing i.e. autophosphorylation; 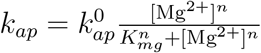

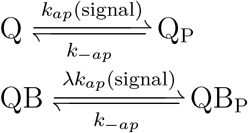
- PhoQ-MgrB binding

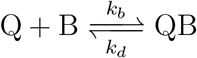
- Phosphotransfer

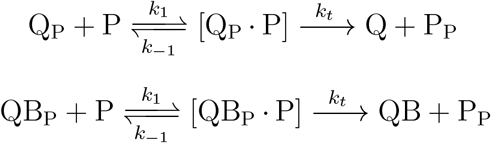
- Dephosphorylation

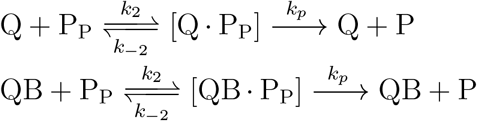

#### Transcription

Modeling of transcription and translation is the same for both models. PhoP-P binds promoter sites as a dimer, therefore we consider the following binding reactions. 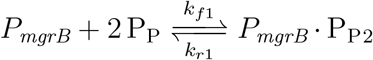

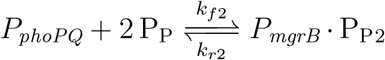

We consider the binding reactions to be quasi-steady state. Considering 1 copy of promoter per cell, the occupancy fraction of the promoter can be calculated as:

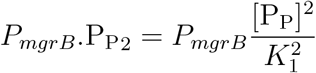

Here 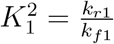.

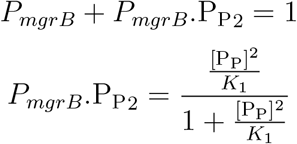

Transcription from a promoter occurs at two rates: basal rate and active rate (when promoter is bound by the transcription factor). All transcripts degrade at a constant rate *k_md_*

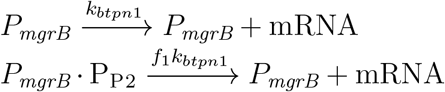

The same calculation gives a model for transcription for any mRNA (PhoP, PhoQ, MgrB or fluorescent reporter) from any PhoP-P dependent promoter (*P_mgrB_* or *P_phoPQ_*).

#### Translation

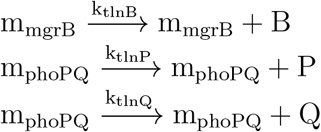

*k_tlnQ_* is assumed 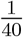 times *k_tlnP_*. All proteins are assumed stable and dilute due to cell growth at a rate *k_pdeg_*.

#### Model ODEs: PhoPQ model with single bifunctional kinase

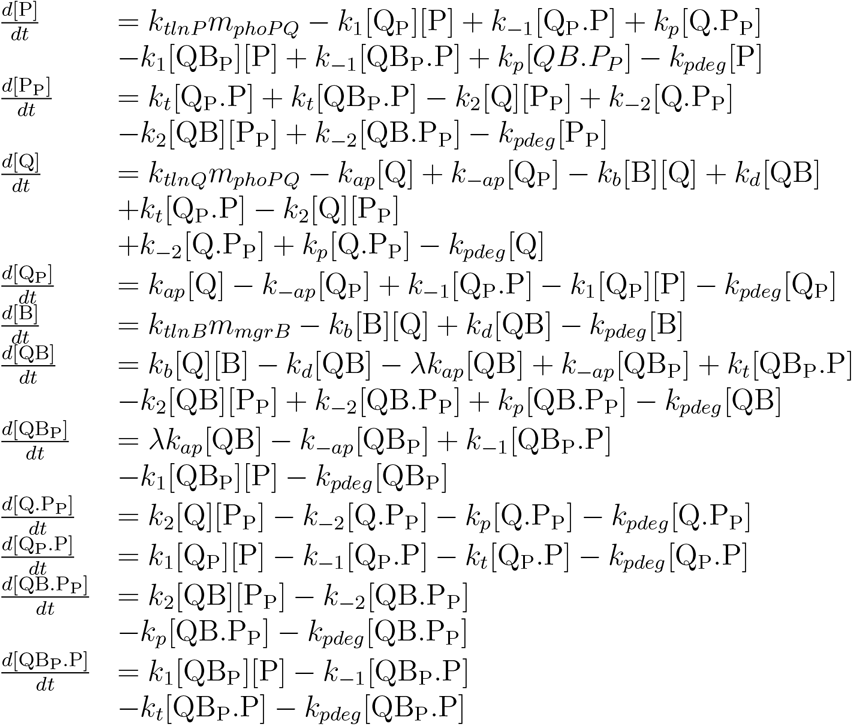

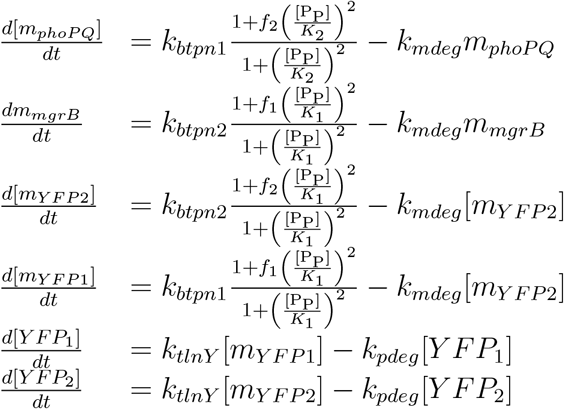

##### *in-silico* mutants

*In-silico* mutants are modeled in the same way for both dynamical models.

1. MgrB-deletion mutant, Δ*mgrB*: Set *k*_*btpn*2_ = 0 in 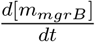 only. This leads to zero MgrB but the reporter mRNA (mYFP2) and reporter protein (YFP) continue to report *P_mgrB_* activity.
2. Autoregulation deletion: Set *f*_1_ = 1 in 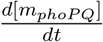 only so that total PhoP, PhoQ remains constant but reporters mRNA (mYFP1) and protein (YFP1) levels continue to report *P_phoPQ_* activity.
3. Constitutive *mgrB*: Set *f*_2_ = 1 and vary *k*_*btpn*2_ in 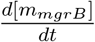 only to set desired total MgrB expression at steady state, while mYGP2 and YFP2 continue to report on *P_mgrB_* activity.
4. PhoQ-phosphatase activity mutant: Set all 3 rates associated with the Michaelis-Menten dephosphorylation reaction zero for Q and QB.
5. Double deletions will be combinations of individual *in-silico* mutants

### Parameters: PhoPQ model with single bifunctional kinase

**Table 1:** Parameters used for one state PhoPQ model simulation (S3 Fig)

| Parameter | Description | Units | Value |
| --- | --- | --- | --- |
| $k_{pdeg}$ | Protein dilution rate | $s^{-1}$ | $3.1 \times 10^{-4}$ |
| $k_{mdeg}$ | mRNA decay rate | $s^{-1}$ | $1.7 \times 10^{-3}$ |
| $k_{tlnP}$ | PhoP translation rate | $s^{-1}$ | $5.0 \times 10^{-2}$ |
| $k_{tlnB}$ | MgrB translation rate | $s^{-1}$ | $4.0 \times 10^{-3}$ |
| $k_{btpn1}$ | basal rate of transcription from $P_{phoPQ}$ | mRNA copies $s^{-1}$ | $4.1 \times 10^{-6}$ |
| $k_{btpn2}$ | basal rate of transcription from $P_{mgrB}$ | mRNA copies $s^{-1}$ | $8.7 \times 10^{-6}$ |
| $f_1$ | Fold increase in transcription rate from $P_{phoPQ}$ | - | 23 |
| $f_2$ | Fold increase in transcription rate from $P_{mgrB}$ | - | 55 |
| $K_1$ | Affinity of PhoP-P to $P_{phoPQ}$ | $\mu M$ | 0.47 |
| $K_2$ | Affinity of PhoP-P to $P_{mgrB}$ | $\mu M$ | 0.45 |
| $k_2$ | PhoQ autophosphorylation rate | $s^{-1}$ | 3.1 |
| $k_{-2}$ | PhoQ-P autodephosphorylation rate | $s^{-1}$ | 0 |
| $k_3$ | PhoQ-P to PhoP association rate | $\mu M^{-1} s^{-1}$ | 0.8 |
| $k_{-3}$ | [PhoQ-P.PhoP] dissociation rate | $s^{-1}$ | $1.2 \times 10^{-2}$ |
| $k_4$ | [PhoQ-P.PhoP] phosphotransfer rate | $s^{-1}$ | $2.1 \times 10^{-1}$ |
| $k_5$ | PhoQ to PhoP-P association rate | $\mu M^{-1} s^{-1}$ | 1.13 |
| $k_{-5}$ | [PhoQ.PhoP-P] dissociation rate | $s^{-1}$ | $9.2 \times 10^{-4}$ |
| $k_6$ | [PhoQ.PhoP-P] dephosphorylation rate | $s^{-1}$ | 2.16 |
| $k_{b1}$ | PhoQ to MgrB association rate | $\mu M^{-1} s^{-1}$ | 0.039 |
| $k_{d1}$ | [PhoQ.MgrB] dissociation rate | $s^{-1}$ | $5.6 \times 10^{-4}$ |

### Model reactions: PhoPQ model with two states of PhoQ

Shorthand - PhoQ = Q, MgrB = B

**Figure 2:**
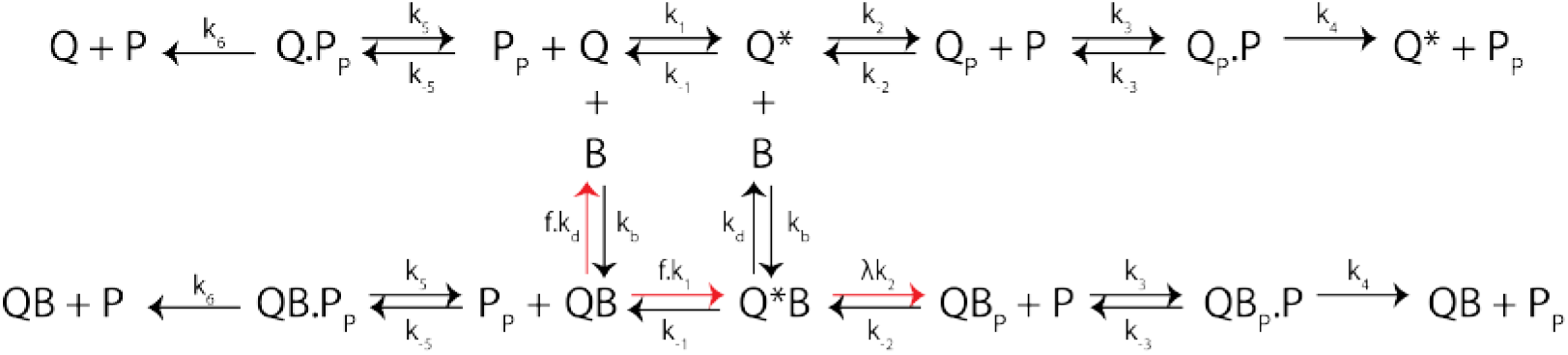
Figure shows all the reactions in the two-state PhoPQ TCS model with associated rate constants. Reactions in the top row represent those in a typical TCS. This modified model includes Q binding B. QB undergoes all the same steps as Q (bottom row) with same rate constants, except autophosphorylation and QB → QB* activation. For detailed balance, we assume QB dissociation is suppressed by the same ratio f.

#### Post-translational reactions

- Signal sensing; 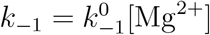

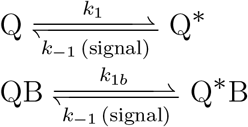
- PhoQ-MgrB binding

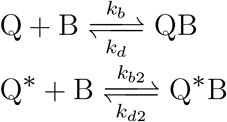
- Autophosphorylation

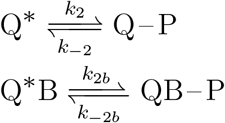
- Phosphotransfer

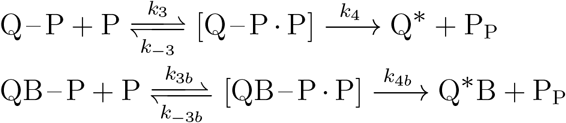
- Dephosphorylation

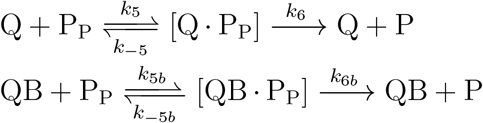

Detailed balance condition constrains parameters as follows:

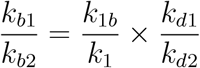

### Model ODEs: PhoPQ model with two states of PhoQ

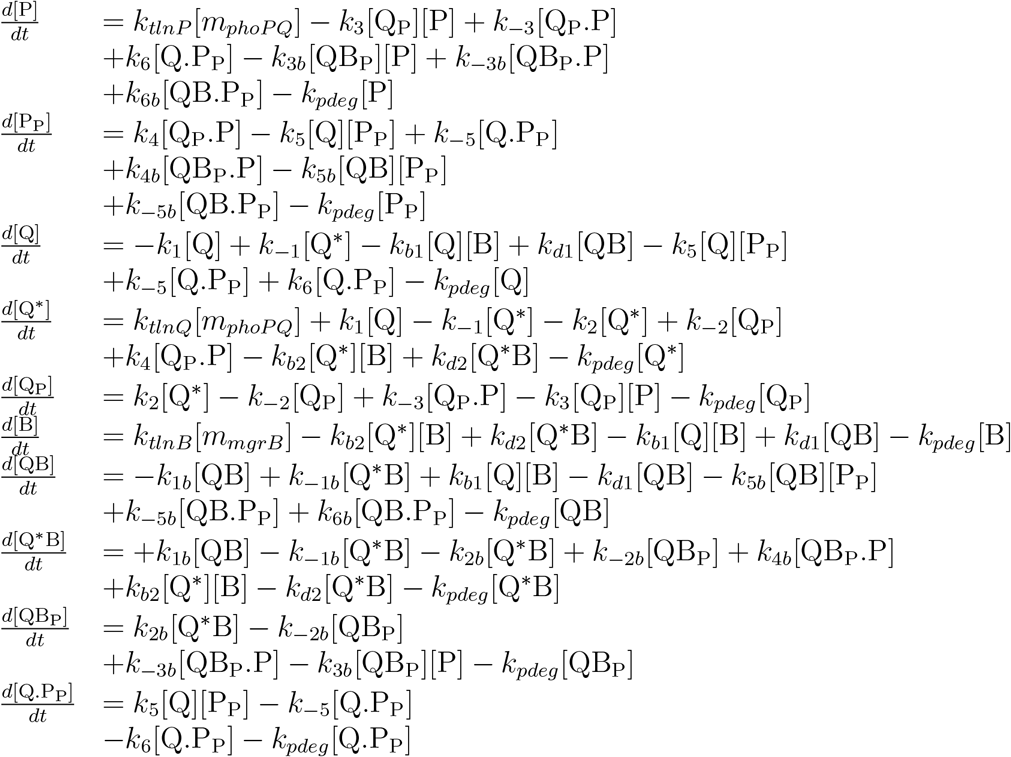

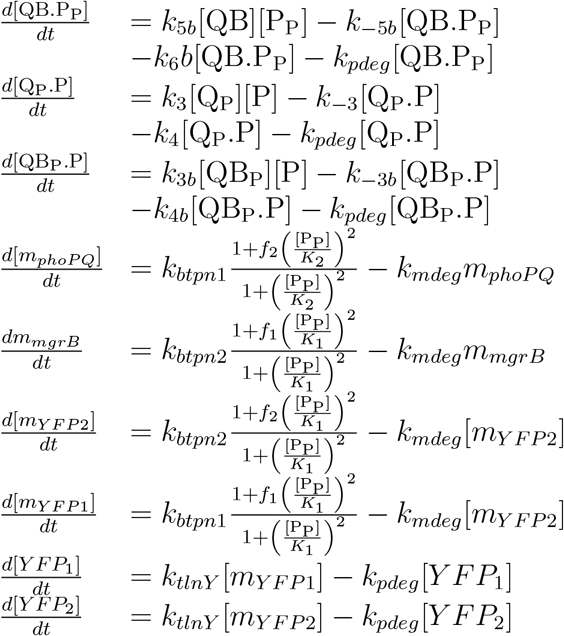

### Parameters: PhoPQ model with two states of PhoQ

**Table 2:**
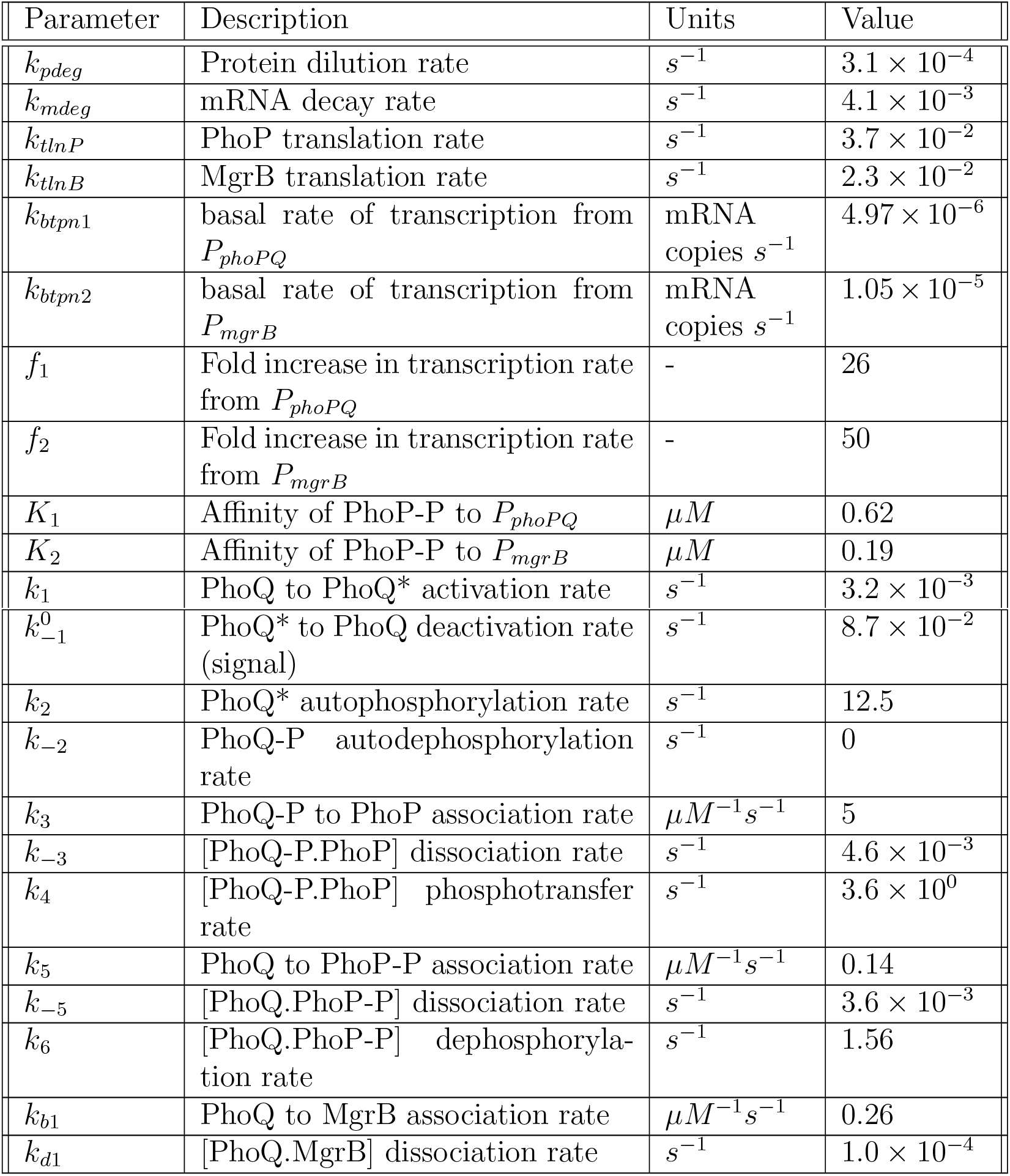
Parameters used for two-state PhoPQ model (S4 Fig)

| Parameter | Description | Units | Value |
| --- | --- | --- | --- |
| $k_{pdeg}$ | Protein dilution rate | $s^{-1}$ | $3.1 \times 10^{-4}$ |
| $k_{mdeg}$ | mRNA decay rate | $s^{-1}$ | $4.1 \times 10^{-3}$ |
| $k_{tlnP}$ | PhoP translation rate | $s^{-1}$ | $3.7 \times 10^{-2}$ |
| $k_{tlnB}$ | MgrB translation rate | $s^{-1}$ | $2.3 \times 10^{-2}$ |
| $k_{btpn1}$ | basal rate of transcription from $P_{phoPQ}$ | mRNA copies $s^{-1}$ | $4.97 \times 10^{-6}$ |
| $k_{btpn2}$ | basal rate of transcription from $P_{mgrB}$ | mRNA copies $s^{-1}$ | $1.05 \times 10^{-5}$ |
| $f_1$ | Fold increase in transcription rate from $P_{phoPQ}$ | - | 26 |
| $f_2$ | Fold increase in transcription rate from $P_{mgrB}$ | - | 50 |
| $K_1$ | Affinity of PhoP-P to $P_{phoPQ}$ | $\mu M$ | 0.62 |
| $K_2$ | Affinity of PhoP-P to $P_{mgrB}$ | $\mu M$ | 0.19 |
| $k_1$ | PhoQ to PhoQ* activation rate | $s^{-1}$ | $3.2 \times 10^{-3}$ |
| $k_{-1}^0$ | PhoQ* to PhoQ deactivation rate (signal) | $s^{-1}$ | $8.7 \times 10^{-2}$ |
| $k_2$ | PhoQ* autophosphorylation rate | $s^{-1}$ | 12.5 |
| $k_{-2}$ | PhoQ-P autodephosphorylation rate | $s^{-1}$ | 0 |
| $k_3$ | PhoQ-P to PhoP association rate | $\mu M^{-1} s^{-1}$ | 5 |
| $k_{-3}$ | [PhoQ-P.PhoP] dissociation rate | $s^{-1}$ | $4.6 \times 10^{-3}$ |
| $k_4$ | [PhoQ-P.PhoP] phosphotransfer rate | $s^{-1}$ | $3.6 \times 10^0$ |
| $k_5$ | PhoQ to PhoP-P association rate | $\mu M^{-1} s^{-1}$ | 0.14 |
| $k_{-5}$ | [PhoQ.PhoP-P] dissociation rate | $s^{-1}$ | $3.6 \times 10^{-3}$ |
| $k_6$ | [PhoQ.PhoP-P] dephosphorylation rate | $s^{-1}$ | 1.56 |
| $k_{b1}$ | PhoQ to MgrB association rate | $\mu M^{-1} s^{-1}$ | 0.26 |
| $k_{d1}$ | [PhoQ.MgrB] dissociation rate | $s^{-1}$ | $1.0 \times 10^{-4}$ |

